# TBX5 drives *Aldh1a2* expression to regulate a RA-Hedgehog-Wnt gene regulatory network coordinating cardiopulmonary development

**DOI:** 10.1101/2021.04.09.439219

**Authors:** Scott A. Rankin, Jeffrey D. Steimle, Xinan H. Yang, Ariel B. Rydeen, Kunal Agarwal, Praneet Chaturvedi, Kohta Ikegami, Michael J. Herriges, Ivan P. Moskowitz, Aaron M. Zorn

## Abstract

The gene regulatory networks that coordinate the development of the cardiac and pulmonary systems are essential for terrestrial life but poorly understood. The T-box transcription factor Tbx5 is critical for both pulmonary specification and heart development, but how these activities are mechanistically integrated remains unclear. We show that *Tbx5* regulates an evolutionarily conserved retinoic acid (RA)-Hedgehog-Wnt signaling cascade coordinating cardiopulmonary development. We demonstrate that Tbx5 directly maintains expression of the RA-synthesizing enzyme *Aldh1a2* in the foregut lateral plate mesoderm via an intronic enhancer that is evolutionarily conserved among terrestrial vertebrates. *Tbx5* promotes posterior second heart field identity in a positive feedback loop with RA, antagonizing a Fgf8-Cyp regulatory module and restricting FGF activity to the anterior. Tbx5/Aldh1a2-dependent RA signaling also directly activates *Shh* transcription in the adjacent foregut endoderm through the conserved MACS1 enhancer. Epithelial Hedgehog then signals back to the mesoderm, where together with Tbx5 it activates expression of *Wnt2/2b* that ultimately induce pulmonary fate in the foregut endoderm. These results provide mechanistic insight into the interrelationship between heart and lung development informing cardiopulmonary evolution and birth defects.

**KEY FINDINGS:**

- Tbx5 regulates second heart field patterning and pulmonary development via retinoic acid (RA) and Hedgehog (Hh) signaling.
- Tbx5 directly maintains transcription of the RA-synthesizing enzyme *Aldh1a2* in the posterior second heart field mesoderm via an evolutionarily conserved intronic enhancer.
- Downstream of Tbx5, RA directly promotes *Shh* transcription through the evolutionarily conserved MACS1 endoderm enhancer.
- Downstream of Tbx5, RA suppresses FGF signaling to pattern the second heart field while promoting a Hedgehog-Wnt2/2b signaling cascade that induces pulmonary fate.

**SUMMARY STATEMENT:** Tbx5-dependent Retinoic Acid signaling regulates an evolutionarily conserved gene regulatory network that coordinates cardiac and pulmonary development.

## INTRODUCTION

Proper integration of the cardiac and pulmonary systems begins during early embryogenesis and is essential for terrestrial life. A key feature of cardiopulmonary development is evolutionarily conserved bi-directional paracrine signaling between the foregut endoderm, which gives rise to pulmonary epithelium, and the cardiogenic mesoderm (Xie et al 2012; Rankin et al 2016; Steimle et al 2018). These signals integrate with lineage specific transcription factors (TFs) in a poorly understood gene regulatory network (GRN). A better understanding of this cardiopulmonary GRN will provide insight into how heart and lung development is orchestrated and inform the molecular basis of life-threatening cardiopulmonary birth defects.

The vertebrate heart forms from two distinct populations of cardiac progenitor cells in the anterior lateral plate mesoderm (lpm), termed the first and second heart fields (FHF and SHF; Kelly et al 2014). The FHF differentiates first and forms the early heart tube, including portions of the two atria and left ventricle. The SHF contributes to the anterior and posterior poles of the developing heart and differentiates later. The anterior SHF (aSHF) is characterized by expression of *Fgf8, Fgf10,* and *Tbx1* and generates the right ventricle, portions of the outflow tract and pharyngeal mesoderm (Rochais et al 2009; Kelly et al 2014). The posterior SHF (pSHF) is characterized by the expression of *Tbx5, Osr1, Foxf1*, (Xie et al 2012; Hoffmann et al 2014; Steimle et al 2018) and generates the atrial septum and sinus venosus. A subset of the pSHF marked by *Isl1, Gli1,* and *Wnt2* expression contains multipotent cardiopulmonary progenitors (CPPs) that give rise to lung mesenchyme, pulmonary vasculature, and myocardium of the inflow tract (Peng et al 2013). CPPs are both the recipient and source of reciprocal signaling with the adjacent pulmonary endoderm essential for heart and lung development.

The T-box TF Tbx5 is a key player in coordinating cardiopulmonary organogenesis; it is cell autonomously required in CPPs for heart development and non-autonomously required for pulmonary development (Xie et al 2012; Hoffman et al 2014; Steimle et al 2018; De Bono et al 2018). In its non-cell-autonomous function Tbx5 is required to activate expression of Hedgehog (Hh) ligands in the adjacent foregut endoderm, which are essential for both heart and lung development (Steimle et al 2018). Endodermal Hh signals back to the lpm stimulating Gli TFs, which cooperate with Tbx5 to directly activate expression of mesodermal *Wnt2/2b* signals that are essential to induce pulmonary fate in the adjacent foregut endoderm (Steimle et al 2018; Goss et al 2009; Harris-Johnson et al 2009). Thus, Tbx5 sits atop a hierarchy establishing the reciprocal mesoderm – endoderm - mesoderm signaling loop that coordinates cardiopulmonary development. A major unanswered question is how Tbx5 non-autonomously activates sonic hedgehog (Shh) ligand expression in the foregut endoderm.

A strong candidate for orchestrating the Tbx5-dependent cell non-autonomous activation of *Shh* in the foregut endoderm is retinoic acid (RA) signaling. RA deficient embryos have multiple cardiac and pulmonary defects (Zaffran et al 2014; Xavier-Neto et al 2015; Perl and Waxman, 2019; Sirbu et al 2020). RA is produced in the lpm by the enzyme Aldh1a2 (Niederreither et al 1999; Metzler and Sandell, 2016) and *Aldh1a2* is co-expressed with *Tbx5* in a subset of the pSHF (Hochreb et al 2003; Ryckebusch et al 2008; DeBono et al 2018). RA regulates *Hh* ligand expression in foregut endoderm (Wang et al 2006; Rankin et al 2016) and patterns the SHF by promoting *Tbx5+/FoxF1+* pSHF identity whilst repressing *Tbx1+/Fgf8+* aSHF fate (Niederreither et al 2001; Sirbu et al 2008; Ryckebusch et al 2008; Deimling et al 2009; Ryckebusch et al 2010; Rydeen and Waxman 2016; Rankin et al 2016; DeBono et al 2018). How the regional production of RA is controlled to pattern the pSHF and regulate *shh* expression remains unknown.

In this study, we tested the hypothesis that Tbx5 coordinates cardiopulmonary development by directly controlling expression of the RA-producing enzyme *Aldh1a2*, and that this RA signal initiates a mesenchyme-epithelial signaling cascade that controls both Hh/Wnt-dependent lung induction and SHF patterning. We show that Tbx5 directly maintains expression of *Aldh1a2* in pSHF via an evolutionarily conserved intronic enhancer. We demonstrate that Tbx5/Aldh1a2-dependent RA signaling regulates anterior-posterior SHF patterning and directly activates *Shh* transcription in the foregut endoderm via an evolutionarily conserved MACS1 endoderm enhancer. Hh/Gli and Tbx5 then cooperate to promote Wnt2/2b expression and lung induction. Our data unify previously unconnected observations in the literature and reveal that Tbx5 regulates an Aldh1a2-dependent RA-Hedgehog-Wnt mesoderm-endoderm-mesoderm signaling network that coordinates pulmonary induction and SHF cardiac patterning.

## RESULTS

### Tbx5 regulates cardiopulmonary development and *Aldh1a2* expression in mouse

To investigate the Tbx5-regulated GRN that coordinates heart and lung development we re-examined our published RNA-seq data of cardiopulmonary (CP) tissue (containing both foregut mesoderm and endoderm) micro-dissected from wild type and *Tbx5^−/−^* mouse embryos at E9.5 (Steimle et al 2018). Differential expression analysis revealed 1,588 up-regulated genes and 1,480 down-regulated genes and in the absence of *Tbx5* (≥1.5 fold change and 5%FDR) (**Fig.1A**; **supplemental table S1**; Steimle et al 2018). Reduced expression of Hh signaling components (*Shh*, *Ihh*, *Patch2*), Hh-targets (*Hhip* and *Gli1*), the lung-inducing *Wnt2/2b* ligands, and pulmonary progenitor marker *Nkx2-1*, indicated a loss of pulmonary fate in *Tbx5^−/−^* mutant CPP tissue (**Fig.1B**). We examined the relationship between pSHF/lung and aSHF/pharyngeal transcripts in *Tbx5^−/−^* embryos by intersecting the Tbx5-regulated transcriptome with gene sets from recent single cell RNA-seq studies of the developing E7.75-E9.5 mouse heart and foregut that define aSHF, pSHF, pharynx, and lung progenitor cells (de Soysa et al 2019; Han et al 2020) (**supplemental tables S2 and S3**). We found that 25% of genes specifically enriched in aSHF or pharyngeal cells (91/366), but only 5% of genes specifically enriched in pSHF or lung progenitors (10/213) from the single cell studies, overlapped with transcripts up-regulated in the *Tbx5^−/−^* mutants (*p< 0.0001, hypergeometric probability test, HGT) (**Fig.1A.**) On the other hand, 34% of the marker genes from the pSHF or lung primordia (72/213) but only 6% of the pSHF+pharynx enriched genes (21/366) were down-regulated in *Tbx5^−/−^* mutants (*p< 0.001, HGT) (**Fig.1A.**) The aSHF-enriched genes up-regulated in *Tbx5^−/−^* CP tissue included well-known patterning genes *Hand1, Irx3, Irx5, Mef2c, Meg3, and Tlx1* as well as FGF signaling components and targets including *Fgf8, Fgf10, Spry1, Spry2, Dusp6* (**Fig1. B**). Gene set enrichment analysis (GSEA) confirmed a statistically significant overrepresentation of aSHF/pharynx genes among upregulated genes (NES=1.58; p<0.0001) and overrepresentation of pSHF/lung transcripts among the downregulated genes (NES= −1.99; p<0.0001) in the *Tbx5^−/−^* CP tissue (**supplemental Fig. S1A,B**). Thus, *Tbx5* mutant embryos exhibit a reduction of the pSHF transcriptional program and gain of aSHF gene expression in the pSHF domain, consistent with recent reports (De Bono et al 2018).

**Figure 1.**
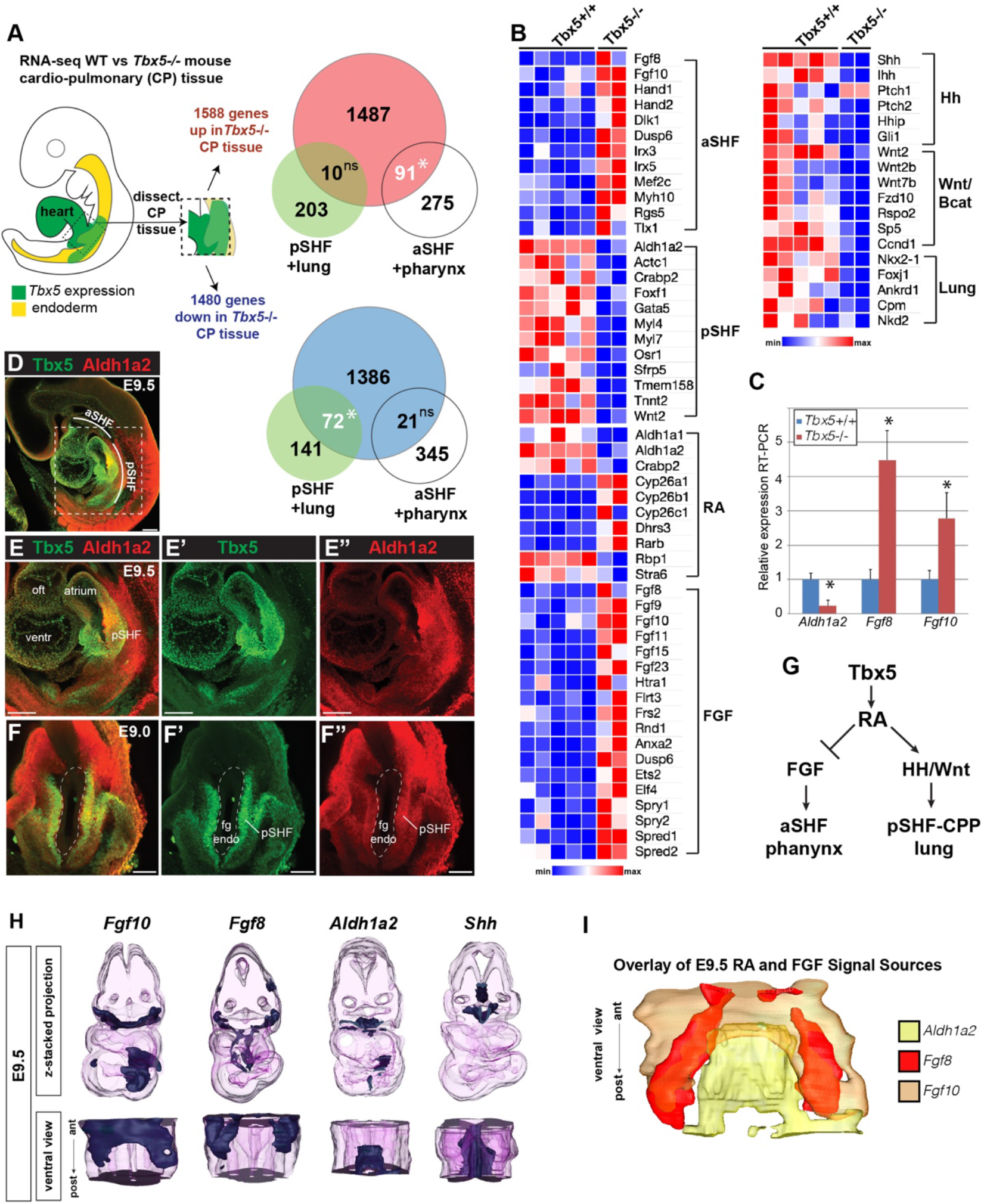
Tbx5 is required for mouse cardiopulmonary development. (A) Schematic of an E9.5 mouse embryo highlighting the dissected cardio-pulmonary (CP) tissue (containing mesoderm and foregut endoderm) profiled by bulk RNA-seq. Venn diagrams show genes differentially expressed in wildtype versus *Tbx5^−/−^* CP tissue (≥1.5 fold change, 5% FDR; Steimle et al 2018; GSE75077) intersected with gene sets from single cell RNA-seq studies defining aSHF + pharynx cells versus pSHF/CPP + lung progenitor cells (de Soysa et al 2019; GSE126128, Han et al 2020; GSE136689) (**supplemental table S2 and S3**). Statistically significant intersection based on hypergeometric tests *p<0.0001. ns= not significant. (B) Transcriptome analysis of Tbx5−/− CP tissue suggests disrupted SHF pattering and failed pulmonary development with reduced RA and increased FGF signaling. Heat map of selected differentially expressed genes in wild-type *Tbx5^+/+^* (n=5) and *Tbx5^−/−^* mutant (n=2) CP tissue grouped by domain of expression or pathway. (C) RT-qPCR validation of decreased *Aldh1a2* and increased *Fgf8, Fgf10* expression in E9.5 wild-type and *Tbx5^−/−^* CP tissue. Relative mean expression + S.D. *p<0.05 Student T-test relative to wild type littermates. (D-F) Wholemount immunostaining of E9.5 (D-E”) or E9 (F-F”) mouse embryos show that Tbx5 (green) and Ald1a2 (red) expression overlaps in a subset of the pSHF. Abbreviations: aSHF=anterior second heart field; pSHF = posterior second heart field; oft = outflow tract; ventr = ventricle; fg endo= foregut endoderm. Scale bar = 100 μM. Model of the proposed Tbx5-RA signaling networks in the cardiopulmonary tissue. 3-D reconstructions from section in-situ hybridizations of e9.5 mouse cardiopulmonary foregut region showing expression domains of *Fgf10, Fgf8, Aldh1a2*, and *Shh* transcripts. Top row is a z-stack 3-D projection; bottom row is a rotated ventral view; (I) shows a ventral view, merged projection of the expression domains of *Fg10, Fgf8*, and *Shh*.

*Tbx5*-dependent genes in the pSHF indicated a possible loss of RA activity. Genes that promote RA signaling were down-regulated in *Tbx5^−/−^* CP tissue, including *Aldh1a2, Crabp2*, which promotes nuclear shuttling of RA, and *Rbp1*, a cytosolic chaperone of the RA precursor retinol (**Fig.1B**). On the other hand, enzymes that attenuate RA signaling, including *Cyp26a1, Cyp26b1, Cyp26c1*, and *Dhrs3*, were increased in *Tbx5^−/−^* pSHF/CPP tissue. Reduced RA-signaling in *Tbx5^−/−^* CP tissue was also consistent with increased expression of multiple TGFβ pathway components and targets (**supplemental Fig. S1C**), which are known to be suppressed by RA during foregut and heart development (Chen et al 2007; Li et al 2010; Ma et al 2016).

The observation that *Aldh1a2* expression was reduced in the *Tbx5^−/−^* pSHF whereas FGF signaling components and targets were increased is consistent with the well-known, evolutionarily conserved role of RA in negatively regulating *Fgf8/Fgf10+* aSHF fate (Rykebusch et al 2008; Sirbu et al 2008; Rydeen et al 2016). RT-qPCR of additional E9.5 dissected CP tissue validated the RNA-seq analysis with *Aldh1a2* being dramatically down-regulated in *Tbx5^−/−^* mutants while *Fgf8* and *Fgf10* were up-regulated **(Fig. 1C.**) Immunostaining of E8.5 - E9.5 wildtype embryos confirmed previous reports that Aldh1a2 is co-expressed with Tbx5 in a subset of the pSHF cells **(Fig. 1D-F and Supplemental Fig. 1SD**) (Hochreb et al 2003; Ryckebusch et al 2008; DeBono et al 2018; de Soysa et al 2019). To investigate the antagonism between RA and FGF signaling within the CP domain, we generated 3D reconstructions using serial sections of *Aldh1a2*, *Fgf8, Fgf10,* and *Shh* in-situ hybridizations through the wild-type mouse E9.5 cardiopulmonary domain. These analyses highlighted the distinct aSHF FGF and pSHF RA signal source domains, with *Aldh1a2+* pSHF mesoderm surrounding the *Shh+* pulmonary foregut endoderm (**Fig. 1H, I**). Together these observations were consistent with the possibility that Tbx5 might regulate SHF patterning by controlling the expression of *Aldh1a2,* which in turn establishes or maintains a local domain of RA activity in the pSHF that subsequently suppresses FGF-mediated aSHF fate and promotes pulmonary development.

### Tbx5 regulates cardiopulmonary development and maintains *aldh1a2* expression in *Xenopus*

To elucidate the underlying molecular mechanisms of the Tbx5-regulated cardiopulmonary GRN, we turned to *Xenopus* embryos, which facilitate the epistatic analysis of signaling pathways. We previously showed that Tbx5-regulated cardiopulmonary development is conserved between *Xenopus* and mouse: Tbx5 loss of function (LOF) in *Xenopus*, either by CRISPR/CAS9-mediated mutation or morpholino (MO) knockdown, phenocopies the mouse *Tbx5^−/−^* phenotype with failed induction of Nkx2-1+ lung progenitors (Steimle et al 2018). Importantly we found that like in mice, *Xenopus tbx5* and *aldh1a2* are co-expressed in the foregut lpm / pSHF and *Xenopus* Tbx5 regulates *aldh1a2* expression (**Fig.2A-C**).

**Figure 2.**
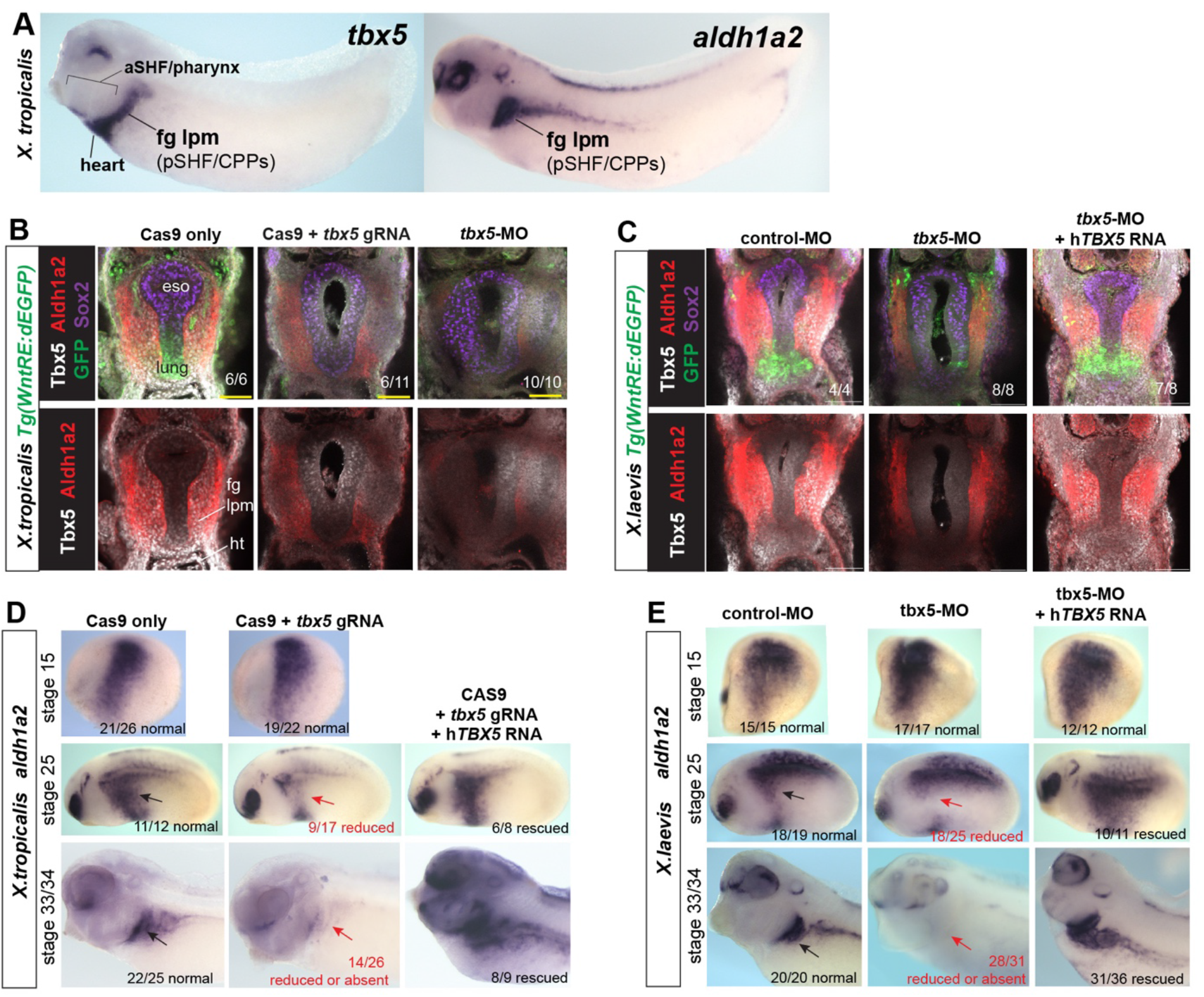
Tbx5 maintains *aldh1a2* expression in *Xenopus* foregut lpm. (A) Whole-mount in-situ hybridization of *tbx5* and *aldh1a2* expression in *X. tropicalis* embryos (NF32), shows co-expression in the foregut lpm that contains pSHF and CPPs. (B-C) Immunofluorescence of transgenic Wnt-reporter embryos *Tg(WntRE:dEGFP)* from (B) *Xenopus tropicalis* or (C) *Xenopus laevis* confirm that like in mouse Tbx5 and Aldh1a2 are co-expressed, and that depletion of Tbx5 by CRISPR/CAS9-mediated mutation or Morpholino (MO) injection results in loss of Aldh1a2 and GFP+ pulmonary fate, which is rescued by co-injection of human *TBX5* RNA. Number of imaged embryos with the observed staining pattern is indicated. Abbreviations: eso, esophagus; ht, heart; pSHF, posterior second heart field. (D-E) Tbx5 is required to maintains *aldh1a2* expression between NF25-34. Wholemount in-situ hybridization of *aldh1a2* expression in *X. tropicalis* (D) or *X. laevis* (E) in control, Tbx5-deficient or Tbx5-rescue embryos at the indicated stage. Expression of *aldh1a2* is unaffected at NF15, reduced by NF25, and largely absent in the fg lpm / pSHF (arrows) at NF33/34. Injection of Human TBX5 RNA (*hTBX5*) rescues *aldh1a2* expression in Tbx5-deficient embryos.

Further analysis showed that Tbx5 LOF in transgenic Wnt/β-catenin reporter embryos *Tg(WntRE:dGFP),* where GFP is under the control of Tcf/Lef sites (Tran et al 2010), resulted in a loss of the GFP in the ventral foregut (**Fig.2B,C**) confirming the failure of Wnt/β-catenin-dependent pulmonary induction. Moreover, cardiopulmonary marker genes that were mis-regulated in the mouse *Tbx5^−/−^* CP tissue were also mis-regulated in *Xenopus* Tbx5 LOF embryos. Pharyngeal/aSHF markers *fgf8*, *fgf10, tbx1, cyp26a1, spry2, hand1, hand2, dhrs3, tgfbR2,* and *tgfbi* were all up-regulated in Tbx5-MO CP foregut tissue, whereas pSHF markers *osr1*, *foxf1*, *gli1* and *wnt2b* and pulmonary endoderm markers *shh, dhh,* and *nkx2-1* were reduced (**Fig.3J**). In total, all 18 transcripts tested exhibited changes in gene expression similar to *Tbx5^−/−^*mice.

**Figure 3.**
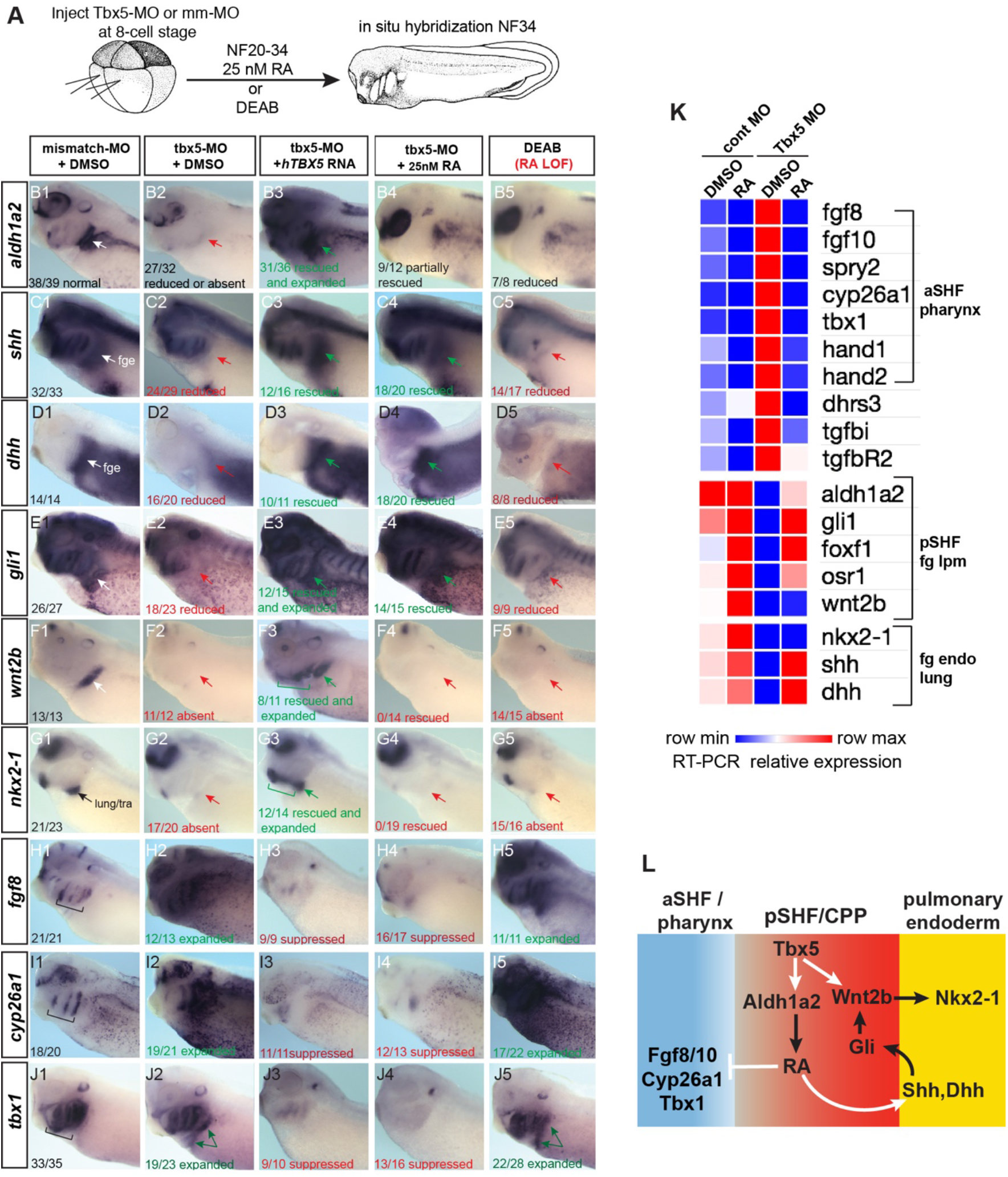
Tbx5 regulates *Xenopus* cardiopulmonary development in part via RA. (A) Schematic of the experimental design. (B-I) Exogenous RA rescues Tbx5 LOF, while inhibition of RA phenocopies Tbx5 LOF. Wholemount in-situ hybridization of NF 34 *X. laevis embryos* after the indicated experimental treatments: injection of negative control 3bp mismatch-MO (10ng), Tbx5-MO (10 ng), Human *TBX5* RNA (*hTBX5;* 100pg), and/or 25nM RA, 10uM DEAB, DMSO vehicle control from NF20-34. The numbers of embryos with the observed expression pattern are indicated. Arrows indicate the relevant expression domain in the CP tissue. Brackets indicate the aSHF/pharyngeal domain. (J) Heatmap showing relative expression from RT-PCR analysis of NF34 Cardiopulmonary (CP)- foregut (fg) tissue dissected from control or Tbx5-MO injected embryos and treated with or without RA from NF20-34. Each row is the average from the 3 biological replicates (n=4 explants per replicate). (K) Diagram of the GRN model showing the key role of Aldh1a2-dependent RA signaling downstream of Tbx5. White arrows indicate relationships tested in the above experiments and black arrows are inferred from previous publications.

Both *X.laevis* Tbx5-MO morphant and *X.trop tbx5* CRISPR/CAS9 mutant embryos exhibited a loss or strong reduction of *aldh1a2* expression in the foregut lpm at NF34 (a timepoint similar to mouse E9.5) (**Fig. 2B-E**). Injection of human *TBX5* RNA rescued *aldh1a2* expression in both *X.laevis* Tbx5 morphants and *X.trop tbx5* mutants (**Fig.2C- E**). A time course analysis revealed that loss of Tbx5 resulted in a downregulation of *aldh1a2* expression in the foregut lpm starting at NF25, but not at early somitogenesis stages (NF15) (**Fig.2D,E**). *Aldh1a2* transcripts were nearly absent and Aldh1a2 protein was strongly reduced in lpm of both Tbx5 morphants and mutants at NF34 (**Fig.2B-E**). These results demonstrate that Tbx5 regulates a conserved transcriptional program in *Xenopus* and mouse to coordinate SHF patterning and lung induction. Moreover *Xenopus* Tbx5 is required to maintain *aldh1a2* expression in the foregut lpm, suggesting that RA signaling is Tbx5-dependent.

### Tbx5 regulates cardiopulmonary development via RA signaling

We tested if failed cardiopulmonary development in Tbx5-deficient *Xenopus* embryos was primarily caused by reduced Aldh1a2-dependent RA signaling. To do so, we asked whether blocking production of endogenous RA could phenocopy loss of Tbx5 or if addition of exogenous RA could rescue the Tbx5 LOF phenotype (**Fig.3A**). Addition of the Aldh enzyme inhibitor DEAB between NF20-34, the time when *aldh1a2* expression was Tbx5-dependent, phenocopied the Tbx5 LOF; pSHF and pulmonary fate markers were reduced/lost and aSHF gene expression was increased/expanded (**Fig.3B-J**). In addition, we treated Tbx5-MO embryos with exogenous RA between NF20-34, using a physiological concentration of 25nM (Horton and Madden 1995; Mic et al 2003; Sheikh et al 2014). As predicted, RA suppressed the expanded expression domains of aSHF markers *fgf8, fgf10, spry2*, *cyp26a1, and tbx1* in Tbx5-deficient embryos (**Fig. 3B-K**). RA was also sufficient to rescue endodermal expression of *shh* and *dhh* as well as expression of known Hh-target genes *gli1, foxf1, and osr1* in foregut lpm of Tbx5-MO embryos and explants (**Fig. 3B-K**). Exogenous RA treatment did not rescue expression of the pulmonary-inducing *wnt2/2b* ligands nor the lung marker *nkx2-1 in* Tbx5 LOF embryos or foregut explants. This observation was consistent with our previous report that Tbx5 is cell autonomously required to directly promote *wnt2/2b* transcription in the pSHF (Steimle et al 2018). Consistent with this interpretation, addition of recombinant WNT2B protein to Tbx5-MO foregut explants was sufficient to rescue *nkx2-1+* lung fate but it did not rescue *shh* nor *dhh* expression (**supplemental Fig.S2**).

These results combined with our previous data suggest that Tbx5 promotes cardiopulmonary development by multiple mechanisms, which are experimentally separable (**Fig. 3L**). First, by maintaining *aldh1a2* expression, Tbx5 ensures robust RA signaling required for proper SHF pattern and for induction of endodermal *Hh* expression. Second, Tbx5 directly activates *Wnt2/2b* expression in the mesoderm (Steimle et al 2018); Hh signals back to the pSHF mesoderm to via Gli TFs, which co-operate with Tbx5 to promote *Wnt2/2b* expression (Rankin et al 2016; Steimle et al 2018), and ultimately Wnt2/2b signal to the foregut endoderm to induce *Nkx2-1+* pulmonary fate.

### Tbx5 directly activates *aldh1a2* transcription and indirectly represses *fgf8* via RA

In preliminary experiments we found that mouse Tbx5 was sufficient to activate *Aldh1a2* transcription and suppress *Fgf8* and *Fgf10* expression during the directed differentiation of mouse embryonic stem cells into cardiac progenitors using a doxycycline inducible *Tbx5* transgenic cell line (Kattman et al 2011; Steimle et al 2018) (**Fig. 4A**). However, in these experiments it was unclear whether Tbx5 regulated *Aldh1a2* or *Fgf* expression directly or indirectly.

**Figure 4.**
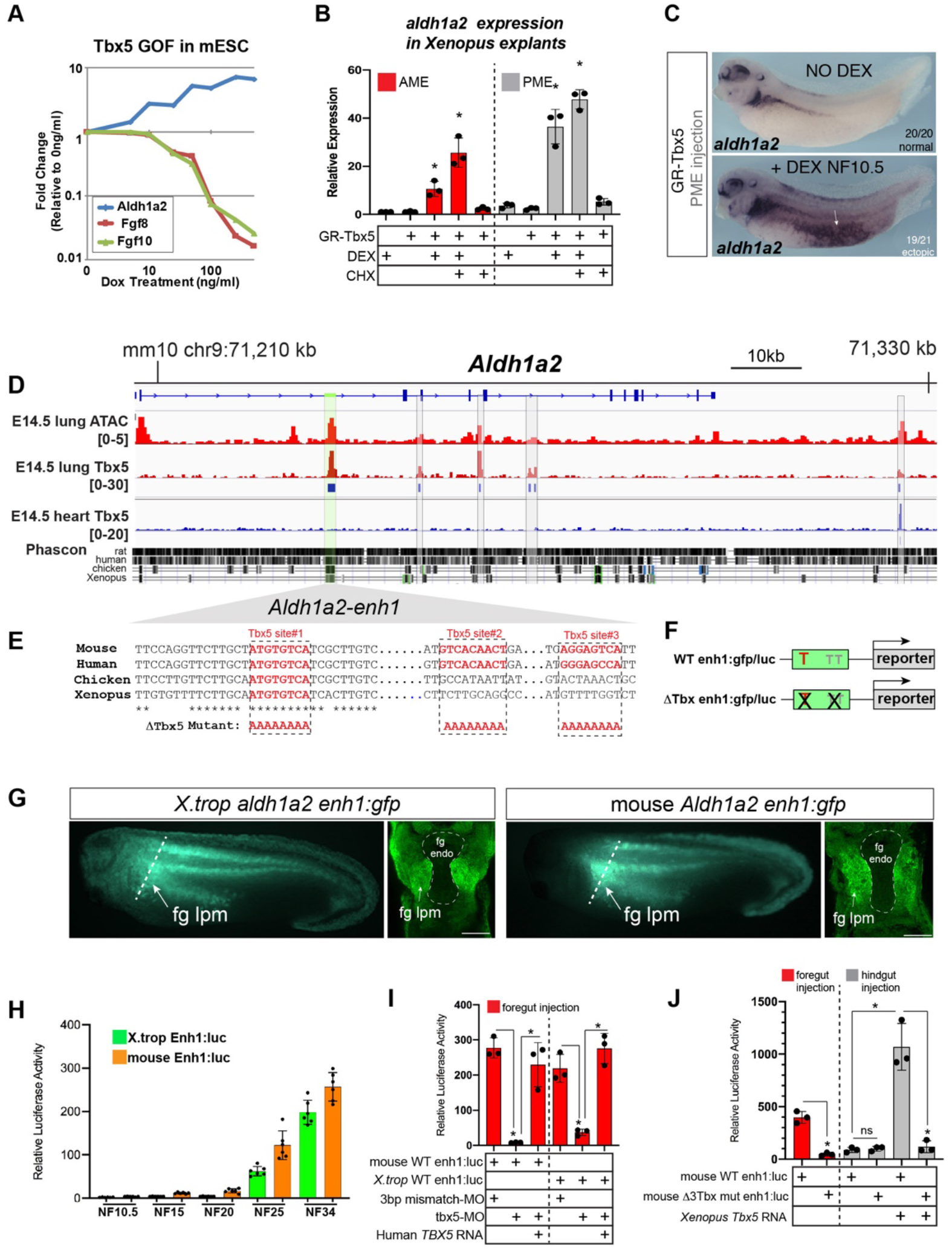
Tbx5 directly activates *aldh1a2* transcription via an evolutionarily conserved first intron enhancer. (A) Dox inducible Tbx5 activated *Aldh1a2* expression and represses *Fgf8* and *Fgf10* mouse embryonic stem cells (mESC) differentiated into cardiac fate in a does dependent manner. Graph shows fold change expression relative to uninduced control. Tbx5 directly activated *aldh1a2* expression in *Xenopus* anterior or posterior mesendoderm (AME, PME) explants. RT-qPCR shows that *aldh1a2* transcription was induced by DEX activated GR-Tbx5 in the presence of CHX. Graphs show mean relative expression ± standard deviation from N=3 biological replicates, each containing 4 explants/replicate. *p<0.05, pair wise Student T-test relative to uninjected, untreated explants. (C) Wholemount in-situ hybridization of *aldh1a2* expression of *X. laevis* NF34 embryos injected with GR-Tbx5 (100pg) into the PME with or without DEX. (D) Genome browser of the mouse *Aldh1a2* locus showing Tbx5 ChIP-seq tracks from E14.5 mouse lung (GSE167207) and E14.5 mouse heart (Burnecka-Turik et al 2020, GSE139803) as well as ATAC-seq track from the ENCODE project (Davis et al 2018; ENCSR335VJW). Tbx5 ChIP-seq peaks in the E14.5 lung are indicated in blue. The multiple species sequence alignment shows that the prominent Tbx5-bound enhancer the first intron (enh1) is evolutionarily conserved from mammals to *Xenopus*. (E) Multiple species sequence alignment of enh1 reveals one Tbx5 DNA-binding site conserved from mammals to *Xenopus* and two additional mammalian-specific Tbx5 sites, which were mutated in reporter constructs. (F) Schematic of the Wild type (WT) and mutant (ΔTbx) enh1:gfp and enh1:luciferase reporter constructs. (G) Both the *Xenopus* and mouse intronic enh1 enhancer are sufficient to drive GFP expression in the foregut lpm in *Xenopus* transgenic assays. Wholemounts show GFP epifluorescence and section views of GFP immunostaining. (H) Time course of *Xenopus* and mouse enh1:luc reporter activity injected into *X*. *laevis* CP-foregut tissue, reflects endogenous Tbx5-dependent *aldh1a2* expression between NF25-34. Graphs show mean relative luciferase activity ± standard deviation. N=5 biological replicates/time point with 5 embryos/replicate. *p<0.05, parametric two-tailed paired T-test. (I) The *Xenopus* and mouse *alldh1a2 enh1* reporter constructs are regulated by Tbx5. Graphs show relative mean luciferase activity ± standard deviation of reporters injected into CP- foregut tissue with control mm-MO, Tbx5-MO, and/or human *TBX5* RNA. N=3 biological replicates/time point with 5 embryos/replicate. *p<0.05, parametric two-tailed paired T- test. (J) The 3 putative Tbx5 motifs in the mouse *alldh1a2-enh1* enhancer are required for reporter activity in the CP-foregut tissue and Tbx5-dependent activation in the hindgut. Graphs show mean relative luciferase activity ± standard deviation. N=5 biological replicates/time point with 5 embryos/replicate. *p<0.05, parametric two-tailed paired T-test.

We therefore examined whether Tbx5 was sufficient to directly activate *aldh1a2* expression in *Xenopus.* We injected RNA encoding a dexamethasone (DEX) inducible Glucocorticoid receptor(GR)-Tbx5 fusion protein (Horb et al 1999) into either the anterior or posterior mesoderm. We then induced GR-Tbx5 nuclear translocation at gastrula stage before endogenous *tbx5* is normally expressed by addition of DEX, with or without the translation inhibitor cycloheximide (CHX) to block secondary protein synthesis (**supplemental Fig.S3A**). GR-Tbx5 activated precocious *aldh1a2* transcription in both the anterior and posterior tissue even in the presence of CHX, demonstrating direct activation (**Fig.4B**). In-situ hybridization of NF34 embryos confirmed robust, ectopic activation of *aldh1a2* by GR-Tbx5 (**Fig.4C**). In contrast, suppression of *fgf8* transcription by GR-Tbx5 was sensitive to CHX, demonstrating indirect repression (**supplemental Fig. S3A,B**). We hypothesized that Tbx5 indirectly represses *fgf8* via Aldh1a2-dependent RA production since RA is known to directly repress *Fgf8* transcription in the mouse SHF (Kumar et al 2016). We tested this by inhibiting Aldh activity with addition of DEAB; this prevented the suppression of *fgf8* by GR-Tbx5 (**supplemental Fig. S3A-C**). Together, these data demonstrate that Tbx5 directly activates *aldh1a2* transcription and indirectly suppresses *fgf8* expression via RA.

### Tbx5 maintains *Aldh1a2* transcription via an evolutionarily conserved intronic enhancer

We next sought to identify *Aldh1a2* enhancers that are directly regulated by Tbx5, predicting that these would be evolutionarily conserved across terrestrial vertebrates. Since a number of putative enhancers have been documented for the mouse *Aldh1a2* locus (Castillo et al 2010; Vitobello et al 2011; Huang et al 2012), we focused on the murine genome. To identify Tbx5-bound enhancers in the CP lineage, we performed Tbx5 chromatin immunoprecipitation followed by high-throughput sequencing (ChIP-seq) of E14.5 fetal mouse lungs as lung mesenchyme is derived from the E9.5 CPPs (Peng et al 2013). ChIP-seq uncovered five Tbx5-bound regions at the *Aldh1a2* locus. Comparing the lung ChIP-seq data to our previously published Tbx5 ChIP-seq from E14.5 heart (Steimle et al. 2018) we found that four of the five Tbx5-bound sites were lung-specific and not bound by Tbx5 in the fetal heart (**Fig. 4D**). Among the four Tbx5-bound sites, only one peak in the *Aldh1a2* first intron, which we refer to as enh1 (for “enhancer 1”; **supplemental Fig.S5A,B**), showed strong evolutionarily conservation from mammals to *Xenopus* (**Fig.4D**). The enh1 region also had a strong ATAC-seq peak in E14.5 lungs indicating open enhancer chromatin (**Fig. 4D**). Sequence analysis of enh1 revealed multiple predicted Tbx5 DNA-binding motifs, one of which was perfectly conserved amongst human, mouse, chicken, and *Xenopus* (**Fig.4E; supplemental Fig.S5A**).

We tested the ability of both the mouse and *X.trop* enh1 intronic enhancers to drive reporter expression in *Xenopus* transgenics and luciferase reporter assays (**Fig.4F-J**). In transgenics, both the mouse and *X.trop* enh1 enhancers drove GFP expression in the foregut lpm at NF34, although the expression domain was broader than endogenous *aldh1a2* (**Fig.4G**), suggesting additional cis-regulatory elements normally refine expression *in vivo*. To quantitatively assess temporal and spatial enhancer activity, we micro-injected enh1 reporters into blastomeres targeting the future CP-foregut or hindgut regions and assayed luciferase activity at a range of developmental stages, from gastrula to tailbud (**Fig.4H; supplemental Fig.S4**). Neither the mouse nor *X.trop* enh1 enhancers drove significant reporter activity during early development at NF 10.5, 15, or 20; however, at NF25 and N34 both the mouse and *X.trop* enh1 enhancers were active in the foregut but not hindgut, (**Fig.4H**; **supplemental Fig.S4A,B**). This timing of enh1 enhancer activity (NF25-34) coincides with the timing at which endogenous *aldh1a2* expression is Tbx5-dependent (**Fig.2**). Together these data demonstrate that the evolutionarily conserved enh1 regulates the temporal and spatial transcription of *aldh1a2* in the foregut lpm/pSHF.

We next tested Tbx5 regulation of the enh1 enhancer by combining reporter assays with Tbx5 loss- or gain-of-function experiments. Tbx5-MO knockdown resulted in a dramatic reduction of the mouse and *X.trop* enh1 reporter activity in CP-foregut tissue at NF34, which was rescued by injection of human *TBX5* RNA (**Fig.4I**). Moreover, injection of *Xenopus* or human *TBX5* RNAs were sufficient to ectopically induce robust enh1 reporter activity in hindgut tissue, which does not express endogenous *tbx5* (**Fig.4J**). Mutation of the single Tbx5-binding site that was perfectly conserved amongst human, mouse, chicken, and *Xenopus* enh1 resulted in a 48% (p=0.0046) and 60% reduction (p=0.0031) of the mouse and frog reporter constructs, respectively, in the foregut and also significantly blunted their responsiveness to ectopic Tbx5 in the hindgut (**supplemental Fig.S4C,D**). Mutation of all three putative Tbx5 motifs conserved amongst mammals in the mouse enh1 (**Fig. 4J**) largely abolished reporter activity in the foregut as well as Tbx5-responsiveness in hindgut injections (**Fig.4J**). We conclude Tbx5 directly maintains *Aldh1a2* expression via multiple T-box motifs found in an evolutionarily conserved first intron enhancer.

### FGF gain-of-function phenocopies Tbx5-loss-of-function in *Xenopus*

In light of the finding that Tbx5-dependent RA signaling suppresses *fgf8* and *fgf10,* we tested if a temporal FGF gain-of-function (GOF) would phenocopy Tbx5 LOF (**Fig.5A**). We treated wild-type CP-foregut explants with recombinant FGF8 protein from NF20-34, the period when exogenous RA was sufficient to rescue Tbx5 LOF. As predicted FGF8 treatment largely phenocopied Tbx5 depletion with increased expression of aSHF/pharyngeal markers *tbx1, fgf10, spry2,* and *cyp26a1, as well as* reduced expression of pSHF and pulmonary endoderm genes *wnt2b*, *shh*, *gli1,* and *nkx2-1* (**Fig.5B)**. We also observed reduced expression of *tbx5* and *aldh1a2* consistent with a feedback loop where FGF restricts Tbx5/Aldh1a2-mediate RA signaling (**Fig.5C**).

**Figure 5.**
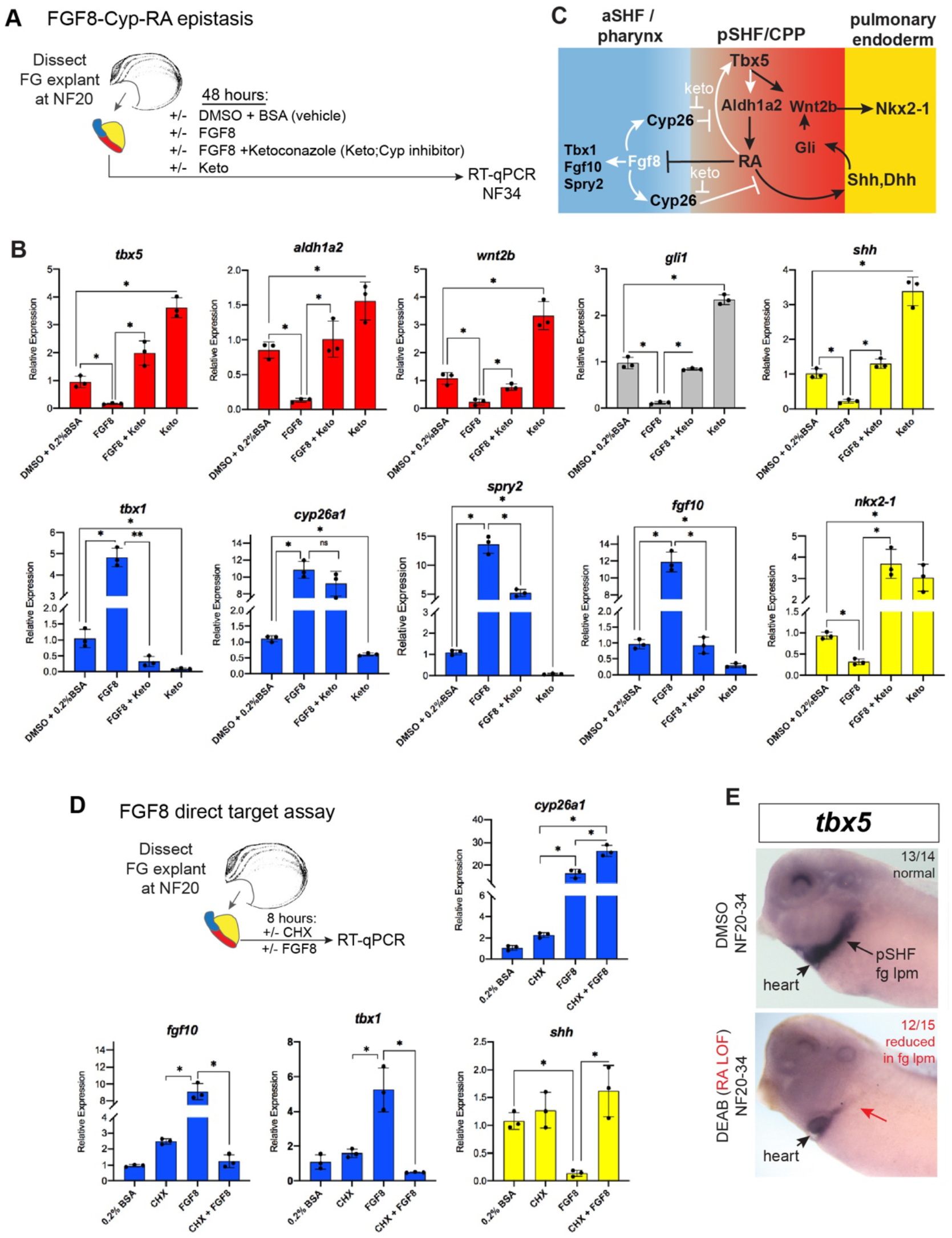
FGF8 gain-of-function phenocopies Tbx5 loss-of-function in *Xenopus*. (A) Schematic of FGF8 gain of function assay in *Xenopus* cardiopulmonary foregut (FG) explants. Explants were micro-dissected at NF20, treated with vehicle controls (DMSO +0.2%BSA) or the indicated combinations of 100 ng/mL FGF8b and/or 0.5 uM ketoconazole (Cyp-inhibitor), harvested at NF34, and analyzed via RT-qPCR. (B) RT-qPCR showing mean relative expression of genes for pSHF (red), aSHF (blue) and pulmonary endoderm (yellow), ± standard deviation from N=3 biological replicates (4 explants/replicate). *p<0.05, parametric two-tailed paired T-test. (C) Model depicting the observed FGF8 gain of function results. White arrows indicate relationships tested in these experiments. (D) FGF8 direct target gene assay in *Xenopus* cardiopulmonary foregut explants, demonstrating FGF8 directly activates *cyp26a1* and indirectly suppresses *shh*. Explants were micro-dissected at NF20, pre-treated with 1uM cycloheximide (CHX) for 2 hours prior to culture in 100ng/mL FGF8b + CHX for 6 hours followed by RT-qPCR analysis. Graphs display mean relative expression ± standard deviation from N=3 biological replicates that contained 4 explants/replicate. *p<0.05, parametric two-tailed paired T-test. (E) RA signaling is required for the *tbx5* expression in the fg lpm / pSHF domain, but not the heart. Embryos were cultured in 10uM DEAB from NF20-34 and assayed by in situ hybridization. Number of embryos assayed and with the observed expression pattern is indicated.

FGF signaling is known to promote the expression of RA-degrading Cyp26 enzymes (Shiotsugu et al 2004; Deimling et al 2011; Rydeen and Waxman 2016), but it is unclear whether this is by direct transcriptional regulation. Therefore, we repeated the FGF8 experiments in the presence of CHX and found that indeed *cyp26a1* was still upregulated by FGF8, demonstrating direct activation (**Fig. 5D**). In contrast, the ability of FGF8 to suppress *shh* was CHX sensitive, demonstrating indirect repression **(Fig. 5D.**) We hypothesized that FGF8 indirectly suppresses expression of *shh* and other RA-dependent pSHF genes by promoting Cyp26-mediated RA degradation (**Fig. 5C**). To test this, we treated CP-foregut explants with both FGF8 and the CYP enzyme inhibitor ketoconazole (keto). Keto blocked the ability of FGF8 to suppress *shh, dhh, tbx5, aldh1a2, wnt2b,* and *nkx2-1* (**Fig. 5B**), indicating that FGF indeed acts via Cyp-dependent RA degradation. Consistent with Cyp-mediated RA degradation being a major factor in endogenous cardiopulmonary patterning, keto treatment alone elevated expression of pSHF (*tbx5, aldh1a2, wnt2b*) and pulmonary endoderm genes (*shh* and *nkx2-1*), whilst decreasing aSHF markers (*fgf8, fgf10, tbx1*) (**Fig.5B**), similar to exogenous RA treatment (**Fig.3**). Interestingly, the CYP inhibition data indicated that RA promotes *tbx5* transcription. Indeed, we verified by in situ hybridization that *tbx5* expression in the pSHF/foregut lpm, but not *tbx5* in the FHF/heart tube, was DEAB-sensitive and required RA (**Fig.5E**). This result is consistent with prior studies in mouse showing RA is required to activate *Tbx5* expression in the pSHF (Niederrereither et al 2001; Ryckebusch et al 2008; De Bono et al 2018). Combined with our finding that Tbx5 directly maintains *aldh1a2* expression, these data identify a RA-Tbx5 positive feedback loop in the pSHF.

### RA directly promotes *Shh* transcription through the evolutionarily conserved MACS1 endoderm enhancer

Our data suggest that RA from the Aldh1a2-expressing lpm is a likely candidate to activate Hh ligand expression in the endoderm. We tested whether exogenous RA could directly activate *shh* and *dhh* transcription in *Xenopus* foregut endoderm explants where the *tbx5/aldh1a2*+ lpm, the source of endogenous RA, had been removed (**Fig.6A,B**). Without the RA-producing lpm, the foregut endoderm did not express *shh* nor *dhh*, however, addition of exogenous RA rescued their expression, even in the presence of CHX, demonstrating direct activation (**Fig.6B)**. As controls RA also rescued expression of the known direct RA-target *hnf1b*, whereas the known indirect target *ptf1a* was not rescued in the presence of CHX (**Fig.6B**).

**Figure 6.**
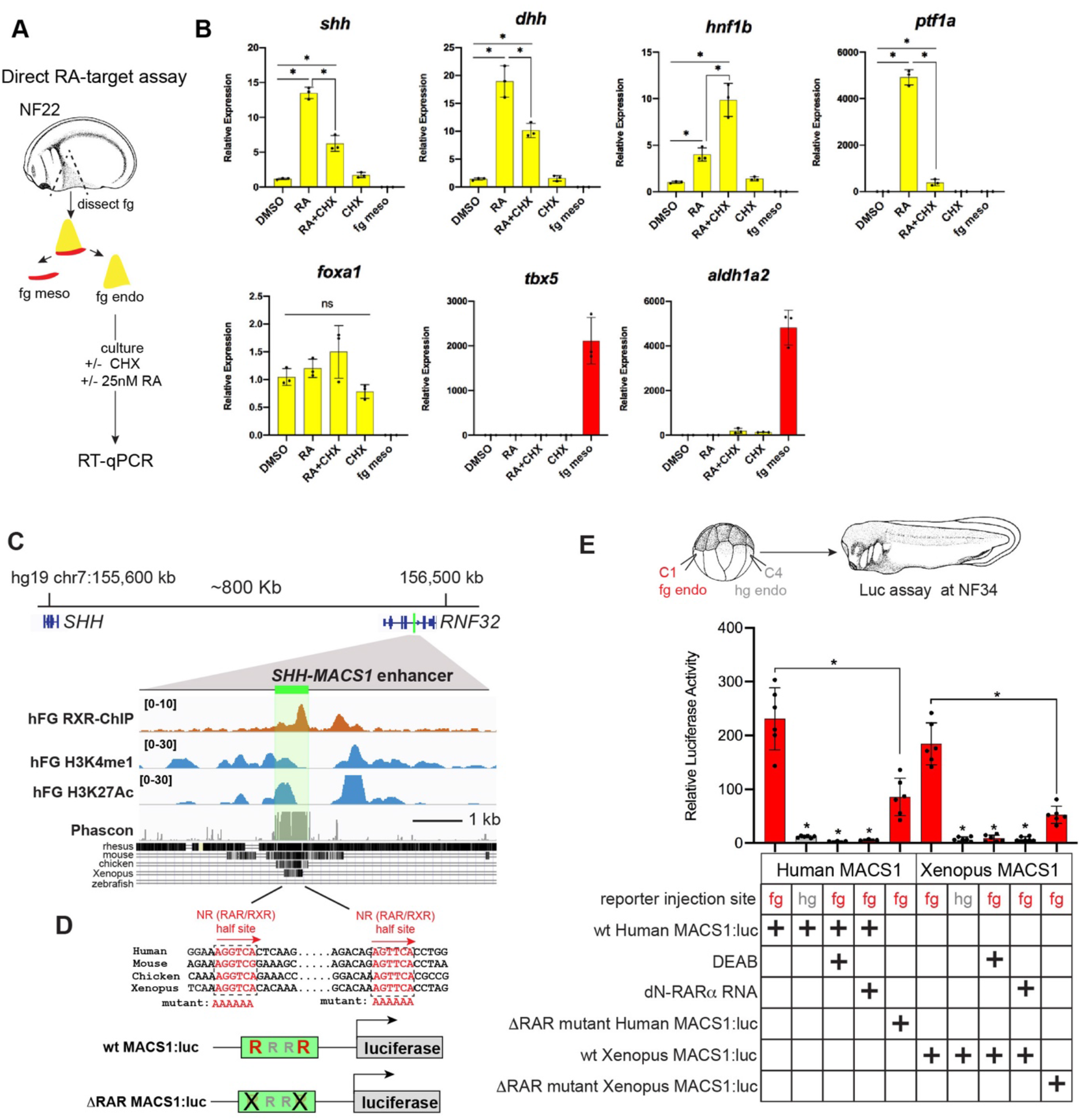
RA-RAR directly activates *shh* transcription in the *Xenopus* foregut endoderm via an evolutionarily conserved MACS1 enhancer. (A) Schematic of direct RA target gene assay in *Xenopus*. Foregut endoderm (fg endo; yellow) was micro-dissected away from foregut mesoderm (fg meso; red) at NF25, pre- treated with 1uM cycloheximide (CHX) for 2 hours prior to culture in 25nM RA + CHX (or DMSO vehicle control) for 6 hours followed by RT-qPCR analysis. (B) RA directly activates *shh* and *dhh* expression in the presence of CHX. Graphs show mean relative expression ± standard deviation from N=3 biological replicates (4 explants/replicate). Endoderm genes are shown in yellow, mesoderm makers in red which confirm clean dissections. *p<0.05, parametric two-tailed paired T-test; ns=not significant. (C) Genome browser of the human *SHH* locus showing the evolutionarily conserved MACS1 distal enhancer known to regulate foregut endoderm expression, which is embedded in an intron of the *RNF32*. Published ChIP-seq tracks of RXR, H3K4me1, and H3K27ac1 from hPSC-derived foregut endoderm (Vinckier et al 2020, GSE104840; Wang et al 2015, GSE54471). Green shading highlights a highly conserved ∼634 basepair MACS1 enhancer region. (D) MACS1 enhancer contains multiple nuclear receptor RAR/RXR DNA-binding half sites, 2 of which are highly conserved. Schematics show the wildtype and mutant MACS1:luciferase reporter constructs. (E) Luciferase reporter assay in *Xenopus* show that the Human and X. *tropicalis* MACS1 enhancers are activated by RA via the RAR/RXR DNA-binding sites. 50pg of MACS1:luciferase reporter+ 5pg pRL-TK reporter were microinjected +/- 250pg of dominant-negative RARa RNA into either the C1 foregut (fg; red bars) or C4 hindgut (hg; grey bars) blastomeres and luciferase activity was assayed at NF34. 10 μM DEAB treatment was from NF20-NF34. Mean relative luciferase activity ± standard deviation, from N=6 biological replicates/time point with 5 embryos/replicate. *p<0.05, parametric two-tailed paired T-test relative to WT MACS1:luc in the foregut (fg).

Previous work has identified an evolutionarily conserved distal *Shh* enhancer called MACS1 (for **<u>m</u>**ammalian-**<u>a</u>**mphibian-**<u>c</u>**onserved **<u>s</u>**equence **<u>1</u>),** which is located more than 800 Kb from *Shh*, within an intron of the *Rnf32* gene (Sagai et al 2009; Tsukiji et al 2014; Sagai et al 2017). The MACS1 enhancer is able to dive transcription in mouse foregut endoderm but the signals and TFs that control *Shh* expression via the MACS1 enhancer are unknown. An analysis of publicly available ChIP-seq data from human foregut endoderm, differentiated from pluripotent stem cells (hPSCs) in part by RA treatment (Vinckier et al 2020; Wang et al 2015), revealed binding of the RA nuclear receptor RXR at the human *SHH* MACS1 enhancer as well as H3K4me1 and H3K27ac1, epigenetic marks indicative of enhancer activation (**Fig.6C**). Sequence analysis of the MACS1 enhancer predicted multiple RXR/RAR nuclear RA receptor half sites (Penvose et al 2019), two of which were evolutionarily conserved between human, mouse, chicken, and *Xenopus* (**Fig.6D; supplemental Fig.S6**). These findings raised the possibility that RA directly activates *SHH* transcription via the MACS1 enhancer.

We functionally interrogated human and *X.tropicalis SHH* MACS1 enhancer activity in *Xenopus* luciferase assays (**Fig.6E**) and found both could drive robust reporter activity in foregut but not hindgut endoderm, demonstrating spatial specificity (**Fig.6E**). Disruption of endogenous RA signaling via DEAB treatment (NF20-34) or injection of dominant-negative RAR alpha RNA (dN-RARa) abolished human and *X.trop* MACS1 enhancer activity (**Fig.6E**). Moreover, mutation of the 2 highly conserved RAR/RXR half sites in the MACS1 enhancers resulted in a dramatic reduction in reporter activity (**Fig.6E**). Together these data indicate that RA signaling directly regulates *shh* expression in foregut endoderm, via conserved RAR/RXR motifs in the conserved *shh* MACS1 enhancer.

## DISCUSSION

### Tbx5 regulates a RA-HH-Wnt GRN that coordinates SHF patterning and pulmonary specification

Our findings reveal a complex and evolutionarily conserved signaling network downstream of Tbx5 that coordinates early development of the cardiac and pulmonary systems (modeled in **Fig.7**). We identify the following aspects of this SHF mesoderm – pulmonary endoderm signaling network: 1) Tbx5 acts in a positive feedback loop with RA, maintaining *aldh1a2* transcription in pSHF mesoderm by direct activation of an *aldh1a2* enhancer; RA is in turn required to maintain *tbx5* expression in the pSHF; 2) downstream of Tbx5, Aldh1a2-dependent RA signaling is necessary to promote pSHF/CPP identity and suppress aSHF fate; 3) Tbx5-RA and FGF-Cyp form mutually antagonistic modules, with Cyp-mediated RA degradation refining the spatial domain of RA activity; and 4) Tbx5/Aldh1a2-dependent RA signaling from the pSHF is required cell non-autonomously to activate endodermal *Hh* ligand expression by direct RXR/RAR activation of the MACS1 enhancer. Ultimately Hh ligands signal back to the pSHF mesoderm to activate Gli TFs, which cooperate with Tbx5 to directly activate *wnt2/2b* transcription; Wnt2/2b then induce pulmonary fate in the foregut endoderm (Rankin et al 2016; Steimle 2018). Thus, during cardiopulmonary development, Tbx5 regulates production of three key paracrine signals, RA, Hh and Wnt. In this way the lung primordia form next to the pSHF-derived atria and pulmonary vessels, in preparation for these two organ systems to be functionally integrated during development.

**Figure 7.**
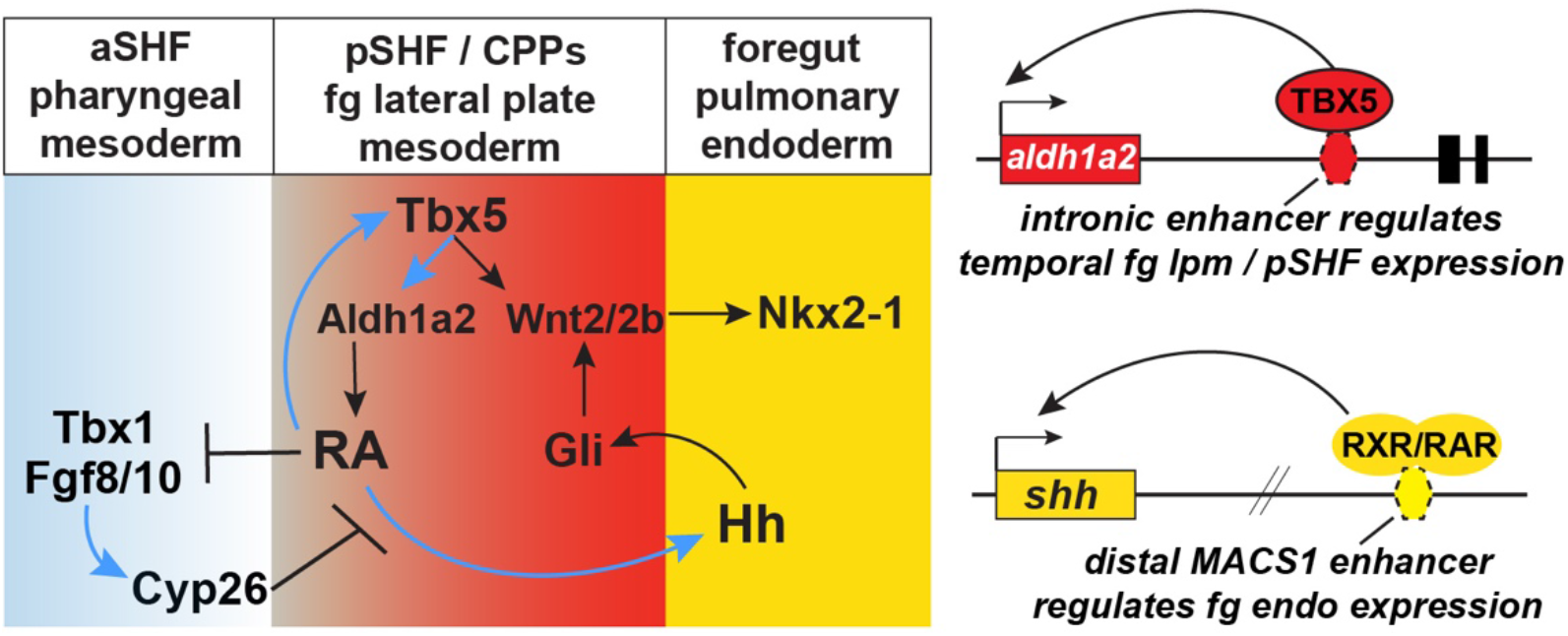
Model of the Tbx5-dependent GRN coordinating SHF pattering and pulmonary induction. Tbx5 directly maintains *Aldh1a2* expression and a RA-Tbx5 positive feedback loop exists in the pSHF, which is necessary for *Hh* ligand expression, Wnt2/2b-dependent pulmonary fate induction, and SHF heart field patterning. Blue arrows in the model indicate relationships tested and demonstrated in this study. Tbx5/Aldh1a2 dependent RA signaling restricts FGF/Cyp activity to the aSHF, promotes pSHF identity, and drives expression of *shh* in pulmonary foregut endoderm. Gene models indicate the locations of the *aldh1a2* enh1 enhancer regulated by Tbx5 and the *shh* MACS1 enhancer regulated by RA/RXR/RAR.

### Enhancers controlling the reciprocal RA-HH mesoderm and endoderm signaling

We identified enhancers driving the direct response of Tbx5 on the *aldh1a2* locus and the direct response of RA at the *shh* locus. Both of these enhancers are highly conserved amongst air breathing terrestrial vertebrate species. We found that Tbx5-dependent RA signaling directly promotes *Shh* expression in foregut endoderm via RAR/RXR half site motifs in the evolutionarily conserved *Shh* MACS1 enhancer (Sagai et al 2009; Tsukiji et al 2014; Sagai et al 2017; Penvose et al 2019). Other endodermal TFs that also contribute to *Shh* expression in the foregut have been described including FoxA2, Meis3, Islet1, Tbx2, and Tbx3 (Lin et al 2006; DiIorio et al 2007; Tamplin et al 2008; Mesbah et al 2012; Lee et al 2019), but whether or not they directly act on the *Shh* MACS1 enhancer, in cooperation with RA/RAR/RXRs, is unknown.

Previous studies have defined multiple enhancers that regulate different temporal and spatial expression domains of *Aldh1a2* during development, with input from T-box TFs as a reoccurring theme. For example, the T-box TFs VegT, Eomesodermin, and Brachyury, regulate *aldh1a2* in the *Xenopus* gastrula mesoderm via cis-regulatory elements that lie upstream of transcription start site (Gentsch et al 2013; Tosic et al 2019). Subsequent expression of *Aldh1a2* in the paraxial mesoderm and early lpm is promoted by Hox/Pbx/Meis TF complexes acting on a first intron enhancer (Vitobello et al 2011) that is distinct from the enh1 enhancer identified herein. We show that Tbx5 directly maintains *aldh1a2* transcription in the pSHF domain of the anterior lpm via the enh1 enhancer. Interestingly Tbx1, which can act as a transcriptional repressor, is also known to spatially restrict *Aldh1a2* expression (Guris et al 2006; Aggarwal et al 2006; Ryckebusch et al 2010), although whether or not Tbx1 directly represses *Aldh1a2* transcription is unclear. We speculate that Tbx1 and Tbx5 might engage the same enh1 enhancer in different cell populations with Tbx1 inhibiting *Aldh1a2* transcription in the aSHF and Tbx5 promoting *Aldh1a2* in the pSHF.

### Tbx-RA feedback loops: a regulatory node in cardiopulmonary birth defects?

We observed that Tbx5 and RA form a positive feedforward loop in the pSHF: in this domain, Tbx5 directly maintains Aldh1a2-dependent RA production and RA maintains *tbx5* expression. Interestingly, RA was not required for *tbx5* expression in the FHF derived heart tube. Conversely, RA signaling is required for *de novo* expression of *Tbx5* in the SHF (de Soysa et al 2019; De Bono et al 2018). We predict that this *de novo* expression in the pSHF is equivalent to the RA-dependent maintenance of *tbx5* that we observed in *Xenopus*. The molecular mechanism by which RA promotes *Tbx5* expression is unclear. Although enhancers that regulate *Tbx5* transcription in the heart and limb have been identified (Minguillon et al 2012; Smemo et al 2012), whether these enhancers control pSHF expression and if they are directly regulated by RA/RAR/RXR complexes remains to be determined.

Interestingly, Tbx1 and RA have an antagonistic relationship during SHF patterning. In mouse, loss of *Tbx1* and aberrant retinoic acid (RA) synthesis result in cardiovascular defects similar to the phenotype found in human DiGeorge syndrome patients. Tbx1 represses *Aldh1a2* expression in the mouse aSHF (Guris et al 2006; Aggarwal et al 2006; Ryckebusch et al 2010) and RA suppresses *tbx1* expression in both *Xenopus* pSHF (our study) and in mice (Ryckebusch et al 2010). Moreover, genetically removing one copy of *Aldh1a2*, thereby reducing the level of RA, decreases cardiovascular malformations found in *Tbx1* heterozygous embryos (Ryckebusch et al 2010). Integrating these Tbx5 and Tbx1 results suggests that T-box TF control of RA signaling acts as a mechanistic toggle, wherein RA is activated by Tbx5 to promote the cardiopulmonary program in the posterior SHF while RA is suppressed by Tbx1 to repress the cardiopulmonary program in the anterior SHF. Tbx5-RA feedforward loops are also required for limb bud development. Limb defects and AVSDs are both a facet of the phenotypic spectrum observed in Holt Oram syndrome in human patients with *TBX5* mutations (Nishimoto et al 2015). This raises the intriguing possibility that Tbx5-RA interactions were an evolutionary innovation in both limb and cardiopulmonary mesoderm in the adaptation to terrestrial life and that disrupting this feedforward loop is central to *TBX5*-associated birth defects. Overall, this work provides a framework for understanding the developmental basis of the human birth defects observed in DiGeorge Syndrome and Holt–Oram Syndrome.

## MATERIALS AND METHODS

### Ethics Statement

Mouse and *Xenopus* experiments were performed according to Institutional Animal Care and Use Committee (IACUC) protocols (University of Chicago protocol 71737; Cincinnati Children’s Hospital protocol 2019-0053).

### Xenopus Methods

#### Xenopus Embryo Injections

Wild-type adult *X.laevis* and *X.tropicalis* frogs were purchased from Nasco (Fort Atkinson, WI). Adult transgenic *X.laevis* and *X.tropicalis* Wnt/B-catenin reporter (*Xla.Tg(WntREs:dEGFP)^Vlemx^*, NXR_0064; and *Xtr.Tg(WntREs:dEGFP)^Vlemx^*, NXR_1094) were purchased from the National *Xenopus* Resource (RRID:SCR_013713). Ovulation, in-vitro fertilization and natural mating, embryo de-jellying, and microinjection was performed as described (Sive et al 2000). Plasmids for *Xenopus* GR-Tbx5 (Addgene 117248) *Xenopus* Tbx5 (Addgene 117247) (Horb et al 1999) and *Xenopus* dominant-negative RARa (Sharp and Goldstein, 1997) were previously described. Human *TBX5* (Horizon Discovery OHS5894-202500411) was gateway sub-cloned from its entry vector pENTR223 into the expression vector pCSf107mT-Gateway-3’myc (Addgene 67617) using clonase (ThermoFisher 11791020) according to manufacturer’s instructions. Linearized plasmid templates were used to make mRNA for injection using the Ambion mMessage mMachine SP6 RNA synthesis kit (ThermoFisher AM1340). Total amounts of injected mRNA were as follows: GR-Tbx5 RNA, 125 pg; dN-RARa, 200pg; human TBX5-myc,100pg. Previously validated Tbx5 translation-blocking morpholinos (Brown et al 2005; Steimle et al 2018) were injected at the eight-cell stage (mixture of 2.5 ng each MO1+2 per dorsal marginal zone (dmz) in *X.laevis*; mixture of 0.5ng each MO1+2 per dmz in *X.tropicalis*). MOs were purchased from GeneTools (Philmath, OR) and were as follows: Tbx5-MO1: 5′TTA GGA AAG TGT CTC TGG TGT TGC C 3′; a negative control Tbx5 mismatch MO1 with three nucleotides mutated: 5′T**<u>C</u>**A G**<u>T</u>**A AAG T**<u>A</u>**T CTC TGG TGT TGC C 3′; Tbx5-MO2: 5′CAT AAG CCT CCT CTG TGT CCG CCA T 3′; Tbx5 3bp mismatch MO2: 5′**<u>T</u>**AT **<u>C</u>**AG **<u>A</u>**CT CCT CTG TGT CCG CCA T 3′ (mismatch bases are indicated in bold, underlined).

For F0 CISPR-mediated indel mutations, a sgRNA targeting *X.trop tbx5* exon 5 (DNA-binding domain) that causes frameshift mutations was synthesized in-vitro as previously described (Steimle et al 2018). This exon5 sgRNA causes approximately 40% of injected embryos to have a phenotype (**supplemental table S4**). Briefly, 2nL of a mixture containing 50pg/nl sgRNA with 0.5ng/nl Cas9 protein (PNA Bio CP01-20) was injected on either side of the sperm entry point at the 1-cell stage (total of 200pg sgRNA and 2ng Cas9 protein per embryo).

For *Xenopus* whole embryo small molecule treatments, embryos were cultured in 0.1X MBS + 50 ug/mL gent with concentrations of 1uM dexamethasone, 25nM RA, or 10uM DEAB (Sigma D86256). In all experiments, corresponding amounts of vehicle controls (DMSO or 0.2% fatty-acid free BSA) were used.

Gastrula or CP-foregut explants (containing both endoderm and lpm) were micro-dissected in 1X MBS + 50 μg/mL gentamycin sulfate (gent; MP Biochemicals 1676045) +/- 10U/mL dispase (Corning Life Sciences 354235;to help remove the lpm) and were cultured in 0.5X MBS + 0.2% fatty acid free BSA (Fisher BP9704) + 50 ug/mL gent with the following concentrations of factors: 1uM dexamethasone (DEX; Sigma D4902); 1uM cycloheximide (CHX; Sigma C4859); 25nM all-trans RA (Sigma R2625); 100 ng/mL WNT2B (R&D systems 3900-WN-025); 1uM DEAB (Sigma D86256); 100ng/mL FGF8b (R&D systems 423-F8-025); 0.5uM ketoconazole (Tocris 1103). In CHX experiments, explants were treated for 2 hours in CHX prior to DEX+CHX treatment for 6 hours.

#### Xenopus RT-qPCR

*Xenopus* explants were dissected from embryos of 2 to 3 separate fertilization/injection experiments, frozen on dry ice in 200uL of TRIzol (ThermoFisher 15596018) and stored at −80°C. RNA was extracted using TRIzol and purified using the Direct-zol RNA miniprep plus kit (ZymoResearch R2070); 500ng RNA was used in cDNA synthesis reactions using Superscript Vilo Mastermix (ThermoFisher 11755050), all according to manufacturer’s instructions. qPCR reactions were carried out using Powerup mastermix (ThermoFisher A25742) on ABI StepOnePlus or QuantStudio3 machines. *Xenopus* RT-qPCR primer sequences are listed in **supplemental table S5.** Relative expression, normalized to ubiquitously expressed *odc*, was determined using the 2^−ΔΔCt^ method. Graphs display the average 2^−ΔΔCt^ value +/- standard deviation. Statistical significance (p<0.05) was determined using parametric two-tailed paired T-test, relative to uninjected, untreated explants. Each black dot in the RT-qPCR graphs represents an independent biological replicate containing 4 explants. Heat map of *Xenopus* RT-qPCR gene expression was generated using Morpheus software (https://software.broadinstitute.org/morpheus/) and shows the average 2^−ΔΔCt^ value from 3 biological replicates for each condition.

#### Xenopus In-situ Hybridization

In-situ hybridization of *Xenopus* embryos was performed as described (Sive et al 2000) with minor modifications. Briefly, embryos were fixed overnight at 4°C in MEMFA (0.1 M MOPS, 2 mM EGTA, 1 mM MgSO4, 3.7% formaldehyde), washed 3×5 minutes in MEMFA without formaldehyde, dehydrated directly into 100% ethanol, washed 5-6 times in 100% ethanol, and stored at −20°C for at least 24 hours. Proteinase K (ThermoFisher AM2548) on day 1 was used at 2 ug/mL for 10 minutes on stage NF15,NF25 embryos and 5 ug/mL on NF34 embryos; hybridization buffer included 0.1%SDS; RNAse A (ThermoFisher 12091021) used at 0.5ug/mL; and anti-DIG-alkaline phosphatase antibody (Sigma 11093274910) used at 1:5,000 in MAB buffer (100 mM Maleic acid, 150 mM NaCl, pH7.5) + 10% heat-inactivated lamb serum (Gibco 16070096) + 2% blocking reagent (Sigma 11096176001). Anti-sense DIG labeled in-situ probes were generated using linearized plasmid cDNA templates with the10X DIG RNA labeling mix (Sigma 11277073910) according to manufacturer’s instructions.

#### Xenopus Immunofluorescence

Embryos were fixed in 100 mM HEPES (pH7.5), 100 mM NaCl, 2.7% methanol-free formaldehyde for 2 hours at room temperature, dehydrated directly into Dent’s post-fixative (80%Methanol / 20% DMSO), washed 5 times in Dent’s, and stored in Dent’s at −20°C for at least 48 hours. Embryos were serially rehydrated (75%, 40%, 25% methanol) into PBS+0.1% TritonX-100 (PBSTr). Embryos were then cut in a transverse plane through the pharynx and posterior to the liver to create a foregut sample using a fine razor blade on a 2% agarose-coated dish in PBSTr. Foreguts were subjected to antigen retrieval in 1X R-universal epitope recovery buffer (Electron Microscopy Sciences 62719-10) for 1 hour at 60-65°C, washed 2 × 10 min in PBSTr, blocked for 1-2 hr in PBSTr + 10% normal donkey serum (Jackson Immunoresearch 017-000-001) + 1% DMSO at room temperature, and incubated overnight at 4°C in this blocking solution + primary antibodies: chicken anti-GFP (Aves GPF-1020; diluted 1:1,000), mouse anti-Sox2 (Abcam ab79351; 1:1,000), rabbit anti-Aldh1a2 (Abcam ab96060; 1:500); goat anti-Tbx5 (Santa Cruz Biotechnology sc-17866, 1:350). Secondary antibodies were donkey anti-Chicken 488, donkey anti-rabbit Cy3, and donkey anti-mouse Cy5, donkey anti-goat 405 (Jackson ImmunoResearch 703-546-155, 711-166-152, 715-175-151, and 705-476-147, respectively; all used at 1:1,000 dilution). After extensive washing in PBSTr, samples were incubated overnight at 4 °C in PBSTr + 0.2% DMSO + secondary antibodies. Samples were again extensively washed in PBSTr, dehydrated into 100% methanol, washed 5 times in 100% methanol, cleared and imaged in Murray’s Clear (2 parts benzyl benzoate,1 part benzyl alcohol) on a metal slide with glass coverslip bottom using a Nikon A1R confocal microscope to obtain optical sections.

#### Xenopus Luciferase Assays

The *Xenopus* tropicalis v9.1 genome on Xenbase.org; RRID:SCR_003280 (Karimi et al 2018) was used to define *Xenopus* enhancers. For *Xenopus* luciferase assays, the sequences of the mouse *Aldh1a2* Enh1 (chr9:71241739-71242765; mm10 genome), *X.trop aldh1a2* enh1 (chr3:89631924-89632943; v9.1/xenTro9 genome), *X.trop shh* MACS1(chr6:9535614-9536245; v9.1/xenTro9 genome), and human *SHH* MACS1 (chr7:156459384-156460049; hg19 genome) enhancers, as well as their respective mutant forms, were commercially synthesized (Genscript USA, Piscataway NJ; or IDT DNA, Coralville, IA) and cloned into the pGL4.23 firefly luc2/miniP vector (Promega E8411). For enh1 enhancer assays, embryos were co-injected with 5 pg of pRL-TK:renilla luciferase plasmid (Promega E2241) + 50 pg of the pGL4.23 luc2/miniP enhancer:luciferase plasmid and the following amounts of MOs or mRNAs into each dorsal marginal zone (dmz) region of 4-8cell embryos: 3.5 ng of tbx5-MO or 3bp mismatch-MO; 62.5 pg GR-Tbx5 RNA; 50pg *Xenopus* Tbx5 RNA; 50pg human TBX5- myc RNA. For hindgut mesendoderm injections, the luciferase reporters were injected into the ventral-posterior marginal zone at the 4-8cell stage +/- Tbx5 RNA. For analysis of Shh MACS1 enhancer activity in endoderm, C1 (foregut) or C4 (hindgut) blastomeres were injected at the 16- or 32-cell stage with 5pg pRL-TK+ 50pg MACS1:luc +/- 100pg dnRARa RNA.

Each biological replicate contained a pool of 5 embryos, obtained from 2 to 3 separate fertilization/injection experiments which were frozen on dry ice in a minimal volume of 0.1xMBS and stored at −80°C. To assay luciferase activity samples were lysed in 100uL of 100 mM TRIS-Cl pH7.5, centrifuged for 10 min at ∼13,000 x g and then 25uL of the clear supernatant lysate was used separately in firefly (Biotium #30085–1) and renilla (Biotium 300821) luciferase assays according to the manufacturer’s instructions. Relative luciferase activity was determined by normalizing firefly to renilla levels for each sample. Graph show the average relative luciferase activity +/- standard deviation with dots showing values of biological replicates. Statistical significance was determined by parametric two-tailed paired T-test, *p<0.05.

#### Xenopus transgenesis

Transgenesis was carried out using the I-SceI meganuclease procedure (Ogino et al 2006; Pan et al 2006). *Xenopus* transgenic plasmids were constructed using the pI- SceI-d2EGFP plasmid backbone (Addgene 32674). First, a fragment containing the mouse or *X.trop* enh1 enhancers upstream of a minimal TATA box promoter (Tran et al 2010) flanked by duplicated copies of the 250bp chick B-globin HS4 insulator (Allen and Weeks, 2005; Rankin et al 2011) was commercially synthesized (Genscript USA) and cloned into the ApaI/XhoI sites of pBluescript II KS+ (Agilent 212207). ApaI/XhoI digestion released this fragment, and it was ligated into ApaI/XhoI digested pI-SceI- d2EGFP plasmid. The meganuclease reaction contained 200ng DNA, 2.5 μl I-SceI enzyme (New England Biolabs R0694S; kept at −80°C and used within 1 month of purchase) in 20 μl total volume and was incubated at 37°C for 30 minutes. 5nl was then injected twice into 1-cell embryos on either side of the sperm entry point (10nl total of meganuclease reaction injected per embryo). We observed 14/102 (∼13%) and 21/183 (11%) GFP+ full transgenic embryos using the mouse and *X.trop* enh1 constructs, respectively, from two independent injection experiments. As a negative control, 0/87 embryos were GFP positive when injected using reactions that omitted the I-SceI enzyme.

### Mouse Methods

#### RNA-seq

RNA-seq of the micro-dissected e9.5 wild-type and *Tbx5^−/−^* pSHF/CPP was previously published and is available on GEO (Steimle et al 2018; GSE75077). Heat maps were generated using Morpheus software (https://software.broadinstitute.org/morpheus/). Columns in the Fig.1 heat map represent biological replicates (*Tbx5^+/+^* wt N=5, *Tbx5^−/−^* N=2), and each column replicate contained n=4 pooled CP dissected regions

#### RT-qPCR, In situ hybridization, and Immunofluorescence

RT-qPCR of dissected, pooled (n=4) mouse e9.5 pSHF/CPP regions was performed as described (Steimle et al 2018), cDNA generated using SuperScript III First-Strand Synthesis SuperMix (ThermoFisher 18080051), and qPCR was performed using PowerUp mastermix (ThermoFisher A25742). Gene-expression levels were normalized by *Gapdh* and RT-qPCR primers are listed in **supplemental table S5**. In situ hybridization on mouse embryos was performed as described (Hoffmann et al. 2009). *Shh* probe was provided by Elizabeth Grove (University of Chicago). Immunofluorescence of mouse embryos was performed as described (Rankin et al 2016) using rabbit anti-Aldh1a2 (Abcam ab96060; 1:300) and goat anti-Tbx5 (Santa Cruz Biotechnology sc-17866, 1:300).

Reconstructions of whole mount in-situ hybridizations were generated using previously published methods (Steimle et al. 2018). In brief images were obtained and pre- processed using Adobe Photoshop CS3 Extended (version 10.0.1, http://www.adobe.com) and reconstructed with AMIRA (version 5.3.2, http://www.amira.com). Manual review of each image in the stack was performed and corrections made when necessary. LabelFields for gene expression and tissue were generated from the same series of sections using separate CastField and LabelVoxel modules. The SurfaceGen module was used to generate surfaces from these LabelFields. Gene expression models for two different genes were initially aligned using the Landmark (2sets) module, and a minimum of three landmarks were used to align the separate models. These landmarks were located using the pharyngeal endoderm and ventral edge of the SHF. Final alignments were fine-tuned manually using the Transform editor.

#### Mouse ESCs

The inducible *Tbx5*OE-mESC line was previously generated (Steimle et al., 2018) and differentiated to the cardiac lineage as described (Kattman et al 2011). Doxycycline (Sigma D9891; concentrations of 0, 5, 10, 25, 50, 100, 250, 500ng/mL) was applied at the cardiac progenitor-like stage (day 6) and cells were harvested for RNA 24 hours later.

#### ChIP-seq

ChIP-seq was performed using dissected whole lungs from E14.5 CD-1 mouse embryos obtained from Charles River. Chromatin was prepared as previously described (Steimle et al., 2018). For immunoprecipitation, the chromatin extract was incubated with 5ug of the anti-TBX5 antibody (Santa Cruz Biotechnology sc-17866; Lot #G1516) at 4°C for >12 hours in a total volume of 200 μL. The immune-complexes were captured by Protein G-conjugated magnetic beads (Life Technologies, 1003D) and washed as previously described (Steimle et al., 2018). The captured chromatin was eluted in ChIP Elution Buffer (10 mM TrisHCl, pH 8.0; 1 mM EDTA; 1% SDS; 250 mM NaCl) at 65°C. After RNase and proteinase K treatment and reverse cross-linking, DNA were purified. High-throughput sequencing libraries from ChIP and input DNA were prepared using NEBNext Ultra DNA Library Prep Kit (New England Biolabs, E7370S). During library preparation, adaptor-ligated DNA fragments of 200-650 bp in size were selected before PCR amplification using Sera-Mag magnetic beads (GE, 6515-2105-050-250). DNA libraries were sequenced using Illumina Hi-seq instruments (single-end 50 base) by the Genomics Core Facility at the University of Chicago.

### Bioinformatics

#### ChIP-seq Analysis

Raw sequencing reads were aligned to the mm10 genome using Bowtie2 (Langmead and Salzberg, 2012) and SAMtools (Li et al., 2009) requiring a minimum mapping quality of 10 (-q 10). Pooled peak calling was performed using default settings of MACS2 callpeak (Zhang et al., 2008) with a q-value set to 0.05 and tag size set to 6 (-q 0.05 -s 6). A fold-enrichment track was generated using MACS2 with the bdgcmp function (-m FE) for visualization on the IVG genome browser (Thorvaldsdottir et al 2013). Public data reanalyzed in this study was downloaded from GEO either as Bigwig files or raw reads which were processed as described above.

#### RNA-seq analysis

RNA-seq of the micro-dissected e9.5 wild-type and *Tbx5^−/−^* pSHF/CPP was previously published and is available on GEO (Steimle et al 2018; GSE75077). This RNA-seq data was re-analyzed using Computational Suite for Bioinformaticians and Biologists (CSBB – v3.0.0) using *ProcessPublicData* module [https://github.com/praneet1988/Computational-Suite-For-Bioinformaticians-and-Biologists]. Differentially expressed genes (DEGs) between *Tbx5^−/−^* and wild-type were identified using RUVSeq, with a threshold of 1.5 fold change and 5% FDR. Expression heat maps were generated using Morpheus (https://software.broadinstitute.org/morpheus/).

DEGs were compared with gene sets from single cell RNA-seq defining aSHF versus pSHF (de Soysa et al 2019; GSE126128) and pharynx versus CPP + lung progenitor cells (Han et al 2020; GSE136689) from the early mouse embryo. We created a cardio- pharyngeal enriched gene set by combining marker genes of aSHF and pharynx mesendoderm, and a cardio-pulmonary gene set by combining markers pSHF, pulmonary mesoderm and lung endoderm. Overlap in gene sets were visualized by Venn diagrams and significant overlaps were defined by hypergeometric tests (HGT). In addition, we assessed the enrichment of upregulated and downregulated DEGs from the *Tbx5−/−* embryos compared to the single cell data sets by gene set enrichment analysis (GSEA) (Subramanian et al 2005).

## ACKNOWLEDGEMENTS

We thank members of the Zorn and Wells labs for suggestions and Lisa Sandell (University of Louisville) for discussions.

## Data availability

E14.5 mouse lung Tbx5 ChIP-seq data generated in this study is available from the Gene Expression Omnibus (GEO) accession number GSE167207.

## Analysis of public datasets

Previously published data that were reanalyzed include: Tbx5-regulated transcriptome, GSE75077 (Steimle et al 2018); single cell RNA-seq defining mouse aSHF and pSHF, GSE126128 (de Soysa et al 2019); single cell RNA-seq defining mouse E9.5 pharynx, CPP, and lung progenitor cells, GSE136689 (Han et al 2020); from the early mouse embryo, E14.5 mouse heart Tbx5 ChIP-seq GSE139803 (Burnicka-Turek et al 2020); E14.5 mouse lung ATAC-seq (https://www.encodeproject.org; Davis et al 2018; ENCSR335VJW); ChIP-seq of human PSC-derived foregut endoderm for RXR, GSE104840 (Vinckier et al 2020); human PSC-derived foregut H3K4me1 and H3K427Ac, GSE54471 (Wang et al 2015).

## SUPPLEMENTAL FIGURE LEGENDS

**Supplemental Figure S1 – relevant to Fig.1.**
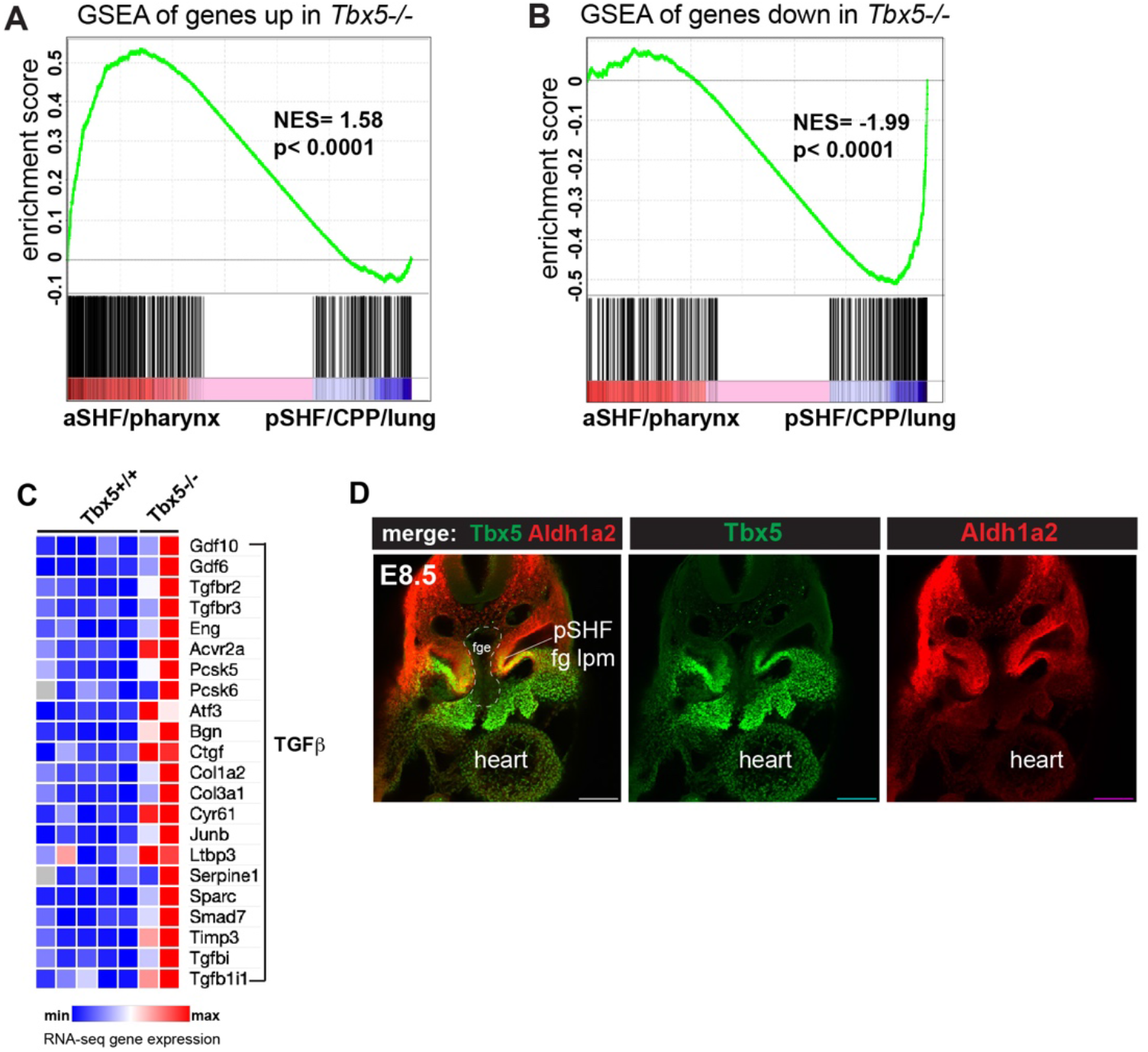
Analysis of the Tbx5 regulated transcriptome in cardiopulmonary tissue. (A-B) Gene set enrichment analysis (GSEA) comparing Tbx5-regulated transcriptome in CP tissue (~≥1.5~ fold change, 5%FDR; Steimle et al 2018, GSE75077) with gene sets from single cell RNA-seq studies defining aSHF versus pSHF (de Soysa et al 2019; GSE126128) and pharynx versus CPP+lung progenitor cells (Han et al 2020; GSE136689) (these gene sets are also listed in **supplemental tables S2**,**S3**). Genes up regulated in *Tbx5^−/−^* CP tissue are enriched in aSHF and pharynx and depleted in pSHF/CPP/lung enriched markers (A). In contrast transcripts that were downregulated in *Tbx5^−/−^* CP tissue were significantly enrichened pSHF/CPP/lung marks and depleted for aSHF/pharynx markers (B). NES= normalized enrichment score. (C) Components and target genes of TGFB signaling are upregulated in *Tbx5−/−* cardiopulmonary tissue. Heatmap of RNA-seq gene expression. Genes listed in the heatmap are known TGFB ligands, receptors, processing enzymes/signaling transducers, and TGFB target genes identified by manual curation of published literature including: Chen et al 2007; Li et al 2010; Ma et al 2016. (D) Immunostaining of e8.5 mouse embryo foregut region shows co-expression of Tbx5 (green) and Aldh1a2 (red) protein in the foregut lateral plate mesoderm (fg lpm) / posterior second heart field (pSHF).

**Supplemental Figure S2 – relevant to Fig. 3.**
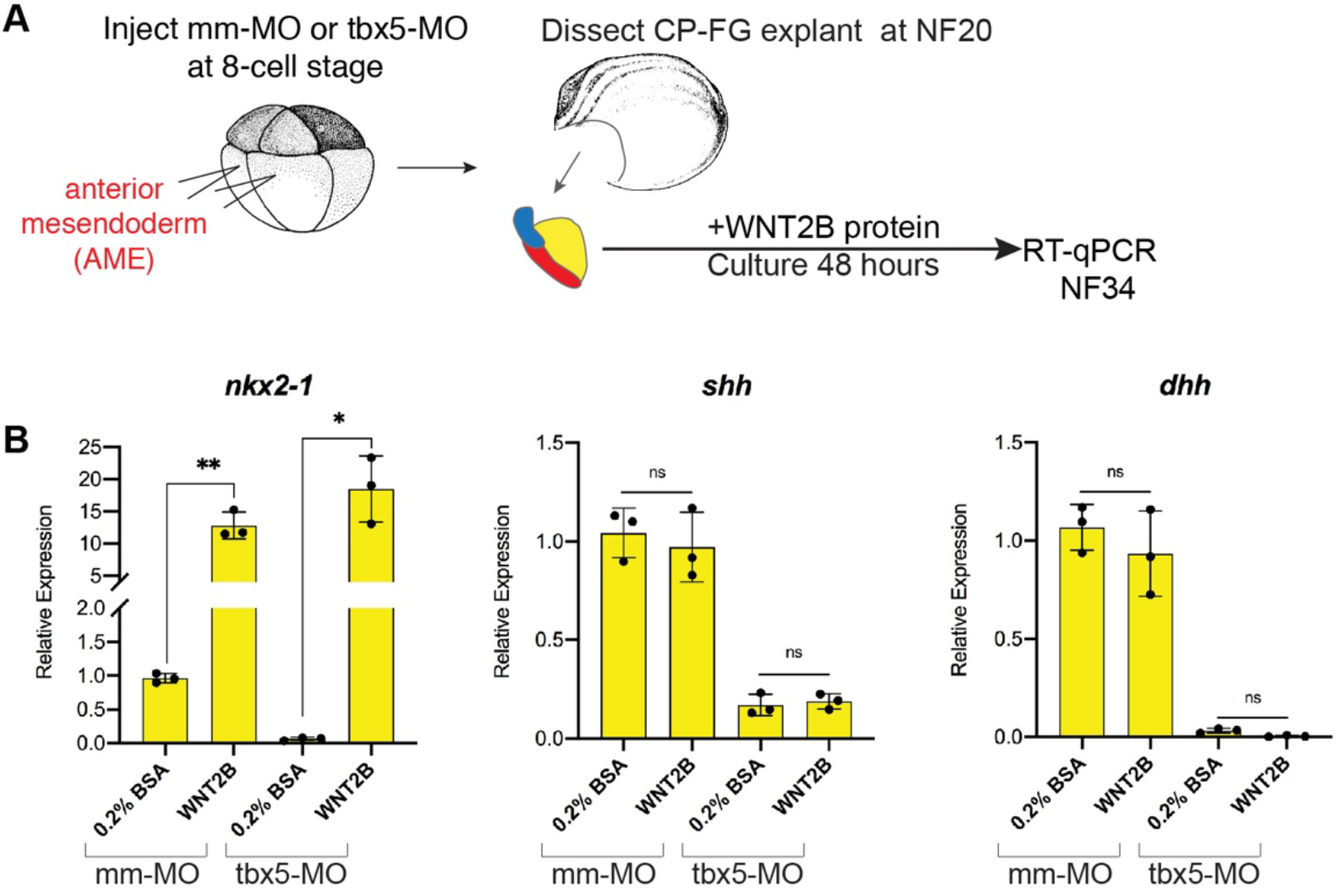
WNT2B protein rescues *nkx2-1+* pulmonary fate in Tbx5-depleted foregut explants. (A-B) WNT2B protein treatment rescues pulmonary *nkx2-1+* fate, but not *shh* or *dhh*, in Tbx5-depleted cardiopulmonary foregut explants. (A) Experimental schematic; (B) 100ng/mL recombinant human WNT2B rescues *nkx2-1* but not *Hh* ligand expression in Tbx5-depleted fg explants. Graphs show mean relative expression ± standard deviation from N=3 biological replicates (4 explants/replicate). Each black dot in the graphs represents a biological replicate (pool of n=4 explants). *p<0.05, **p<0.01, parametric two-tailed paired T-test relative to uninjected, untreated explants; ns= not significant.

**Supplemental Figure S3 – relevant to Fig. 4.**
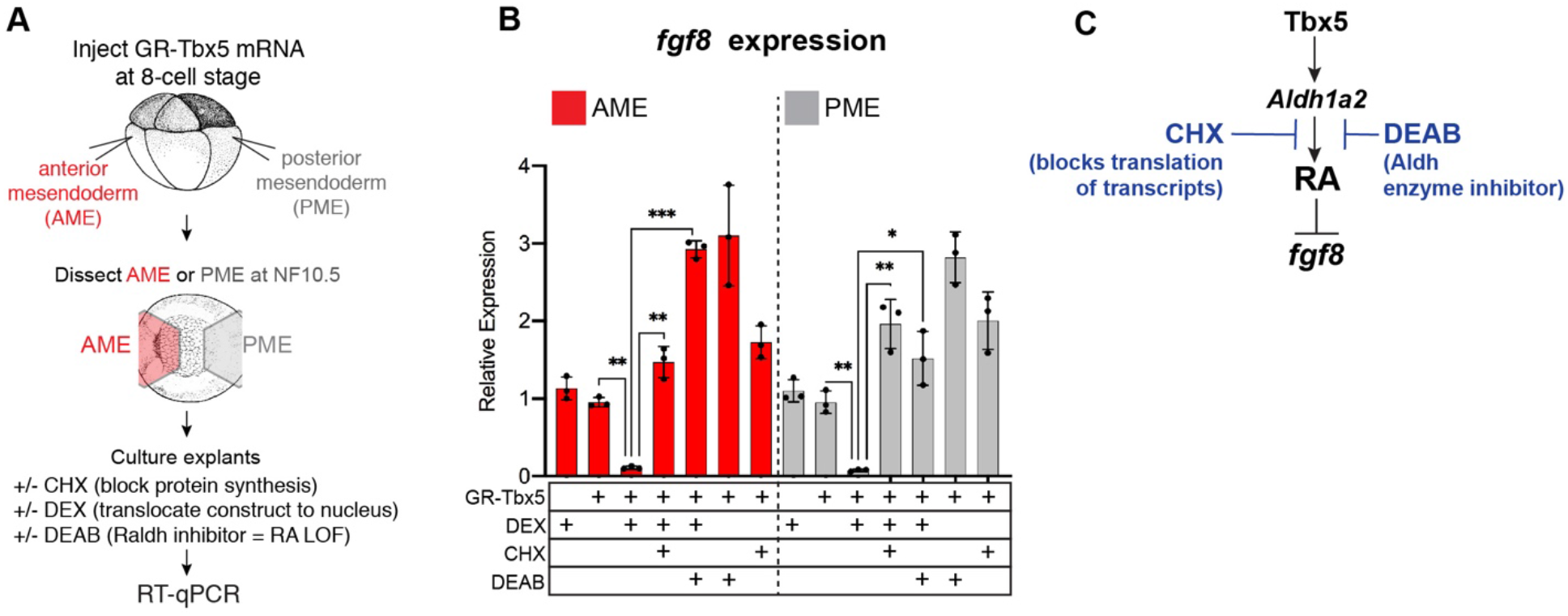
GR-Tbx5 indirectly suppresses *fg8* in *Xenopus* via RA. (A-C) Experimental schematic of GR-Tbx5 direct/indirect target gene assay in *Xenopus* gastrula explants. (B) GR-Tbx5 suppresses fg8 in both anterior and posterior tissue in a CHX sensitive manner demonstrating indirect suppression; the ability of GR-Tbx5 to suppress fgf8 requires is DEAB sensitive and thus is dependent on RA production. Graphs show mean relative expression ± standard deviation from N=3 biological replicates (4 explants/replicate). Each black dot in the graphs represents a biological replicate (pool of n=4 explants). *p<0.05, **p<0.01, p<0.001, parametric two-tailed paired T-test relative to uninjected, untreated explants; ns= not significant. (C) Model of the indirect regulation of *fgf8* by Tbx5, which these experiments show is CHX and DEAB sensitive.

**Supplemental Figure S4 – relevant to Fig. 4.**
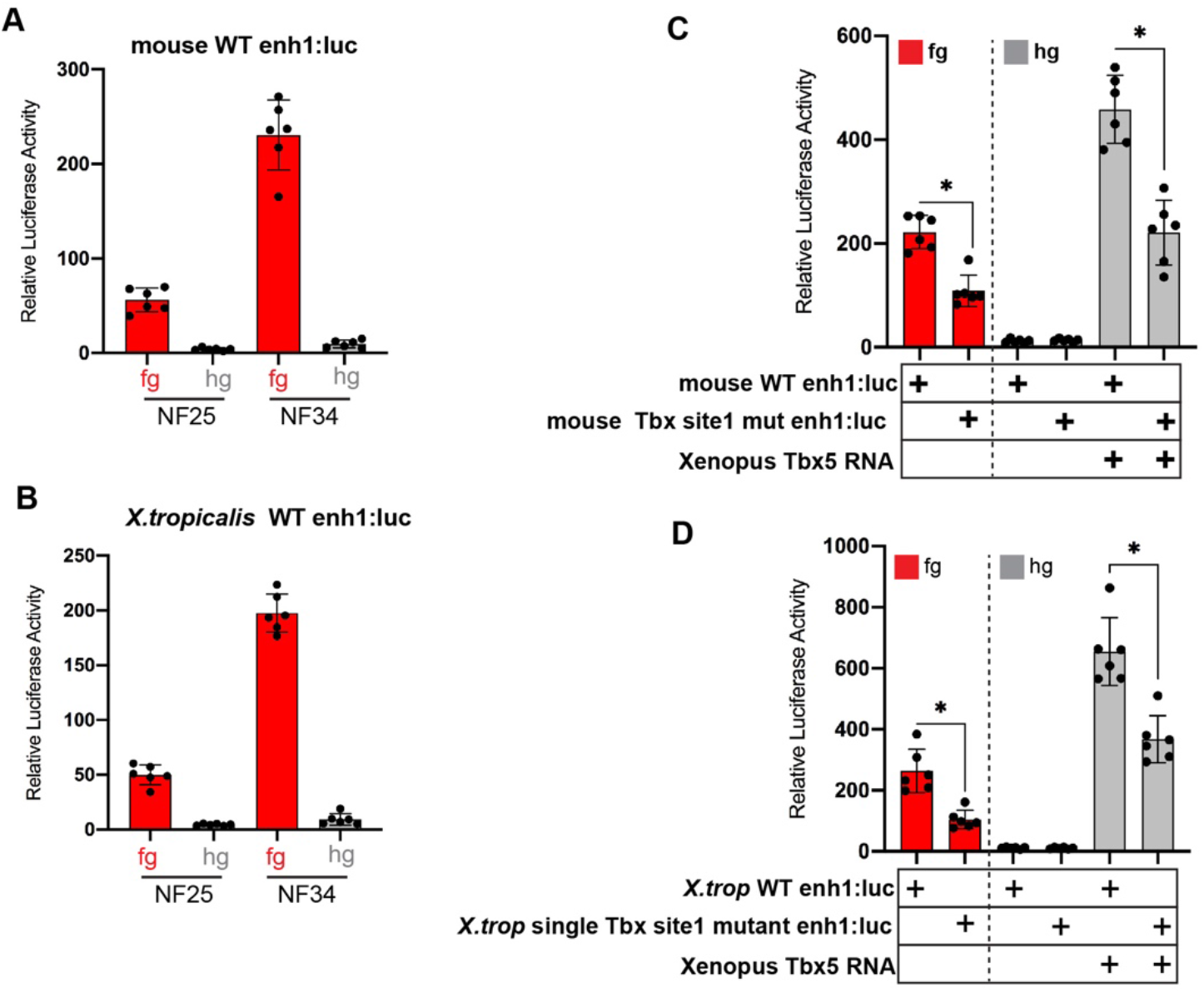
Additional analyses of the mouse and *X.tropicalis Aldh1a2* enhancers in luciferase reporter assays. (A-B) Both the mouse and *X.tropicalis (X.trop) Aldh1a2* enh1 enhancers drive reporter activity in NF25 and NF34 foregut but not hindgut tissue, demonstrating spatial specificity. Graphs show mean relative luciferase activity ± standard deviation. N=6 biological replicates/time point, each containing n=5 embryos/replicate. *p<0.05, parametric two-tailed paired T-test. (C-D) Analysis of wild-type enh1 or enh1 with the single, perfectly conserved T-box/Tbx5 motif mutated. (C) Mutation of the single T-box/Tbx5 motif in the mouse (C) or *X.trop* (D) *Aldh1a2* enh1 enhancers significantly reduces enhancer ability to drive expression in the foregut as well as to be activated by exogenous Tbx5 in hindgut gain-of-function injections. Graphs show mean relative luciferase activity ± standard deviation. N=6 biological replicates/time point, each containing n=5 embryos/replicate. *p<0.05, parametric two-tailed paired T-test.

**Supplemental Figure S5 – relevant to Fig. 4.**
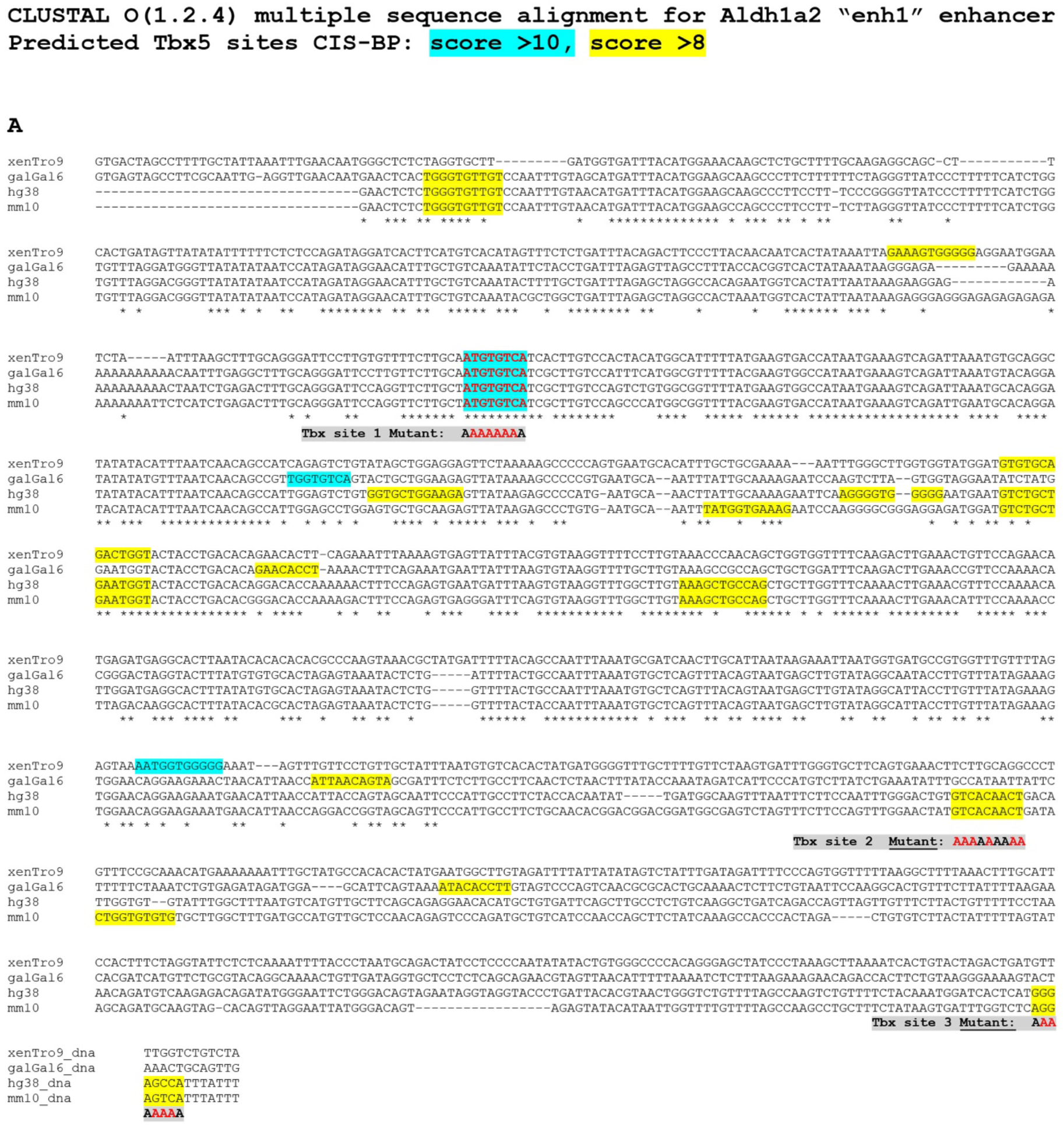
Multiple species sequence alignment of the *Aldh1a2* enh1 enhancers. (A) Clustal DNA sequence alignment; putative T-box/Tbx5 motifs predicted by the CisBP tool (Weirauch et al 2013) are shaded in aqua blue (CisBP score >10) and in yellow (CisBP score >8). Asterisks below the nucleotide alignment indicated conserved bases amongst all four species. Tbx motifs mutated and tested in this study are indicated in gray. Genomic co-ordinates of the *Aldh1a2* enh1 enhancers used were: *X.tropicalis,* >xenTrov9.1_dna range=chr3:89631924-89632943; Chicken, >galGal6_dna range=chr10:7578306-7579190; Mouse, >mm10_dna range=chr9:71241739-71242765 Human, >hg38_dna range=chr15:58038775-58039647

**Supplemental Figure S6 – relevant to Fig. 6.**
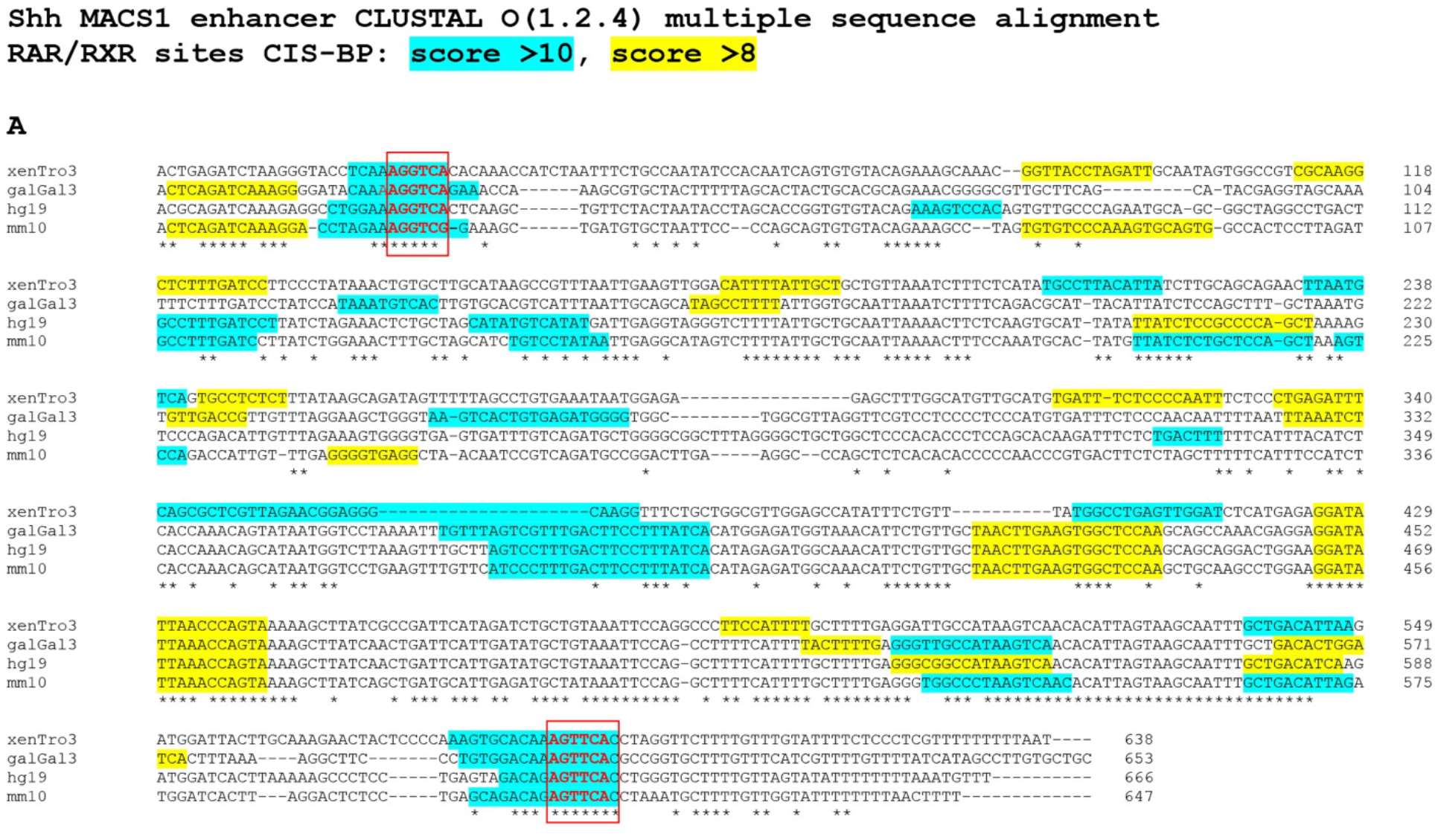
Multiple species alignment of the *Shh* MACS1 enhancers. (A) Clustal DNA sequence alignment; putative RAR/RXR nuclear receptor motifs predicted by the CisBP tool (Weirauch et al 2013) are shaded in aqua blue (CisBP score >10) and in yellow (CisBP score >8). Asterisks below the nucleotide alignment indicated conserved bases amongst all four species. RAR/RXR motifs mutated and tested in this study are indicated and boxed in red. Genomic co-ordinates of the *Shh* MACS1 (mammalian-amphibian-conserved sequence 1; Sagai et al 2009) enhancers used in the multiple alignment were: *X.tropicalis*: v9.1/xenTro9 genome build, chr6:9535614-9536245 Chick: galGal3 genome build, chr2:8370257-8370909 Human: hg19 genome build, chr7:156459384-156460049 Mouse: mm9 genome build, chr5:29538631-29539277

